# A comparative life cycle analysis of electromicrobial production systems

**DOI:** 10.1101/2021.07.01.450744

**Authors:** Anthony J. Abel, Jeremy D. Adams, Douglas S. Clark

## Abstract

Electromicrobial production (EMP) processes represent an attractive strategy for the capture and conversion of CO_2_ into carbon-based products. We describe the development and application of comprehensive reactor, process, and life cycle impact models to analyze three major EMP systems relying on formate, H_2_, and acetate as intermediate molecules. Our results demonstrate that EMP systems can achieve a smaller carbon footprint than traditional bioprocessing strategies provided the electric grid is composed of >~90% renewable energy sources. For each of the three products we consider (biomass, enzymes, and lactic acid), the H_2_-mediated Knallgas bacteria system achieves the lowest overall global warming potential, indicating that this EMP strategy may be best-suited for industrial efforts based on current technology. We also identify environmental hotspots and process limitations that are key sites for future engineering and research efforts for each EMP system. Our analysis demonstrates the utility of an integrated bioelectrochemical model/life cycle assessment framework in both analyzing and aiding the ecodesign of electromicrobial processes and should help guide the design of working, scalable, and sustainable systems.

## Introduction

Ongoing and worsening ecological and humanitarian crises caused by anthropogenic climate change have precipitated efforts to transition away from fossil fuel-based commodity chemical production. Whole-cell biocatalysis provides a theoretically carbon neutral method of producing value-added products if all of the required carbon is originally fixed from atmospheric carbon dioxide (CO_2_). Many petroleum-based products including fuels, plastics, and commodity chemicals can be produced biologically.^1–3^ Moreover, some products, such as proteins, can only be produced biologically and have wide-ranging applications including in food production, chemical sensing, and as therapeutics.^4–6^ Traditional bioprocesses rely on heterotrophic microbes that require exogenous sources of carbon and energy (Fig. 1).

**Figure 1.**
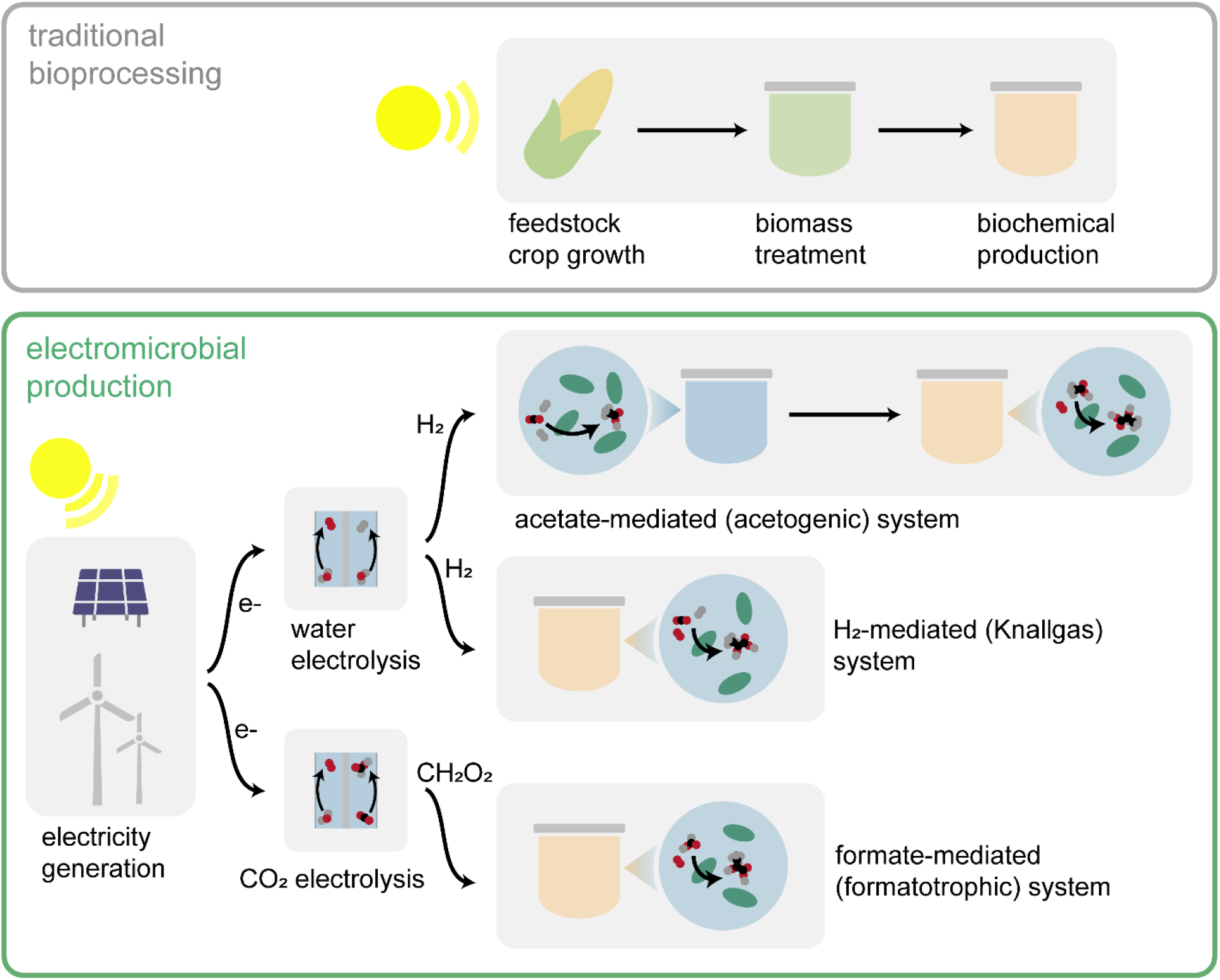
Overview of traditional bioprocessing and electromicrobial production. Traditional bioprocessing relies on feedstock crop growth, pretreatment of the resulting biomass (typically enzymatic or chemical), and subsequent biochemical production using crop-derived sugars as the feedstock. Electromicrobial production uses electricity (ideally renewable) to produce energy substrates (*e.g*., H_2_) for biochemical production from CO_2_.

Glucose from corn starch and sucrose from sugarcane are currently the most common feedstocks in bioprocessing. These biochemical processes rely on extensive agricultural production and therefore compete with the food supply and require land use changes that have significant negative impacts on the environment. Moreover, the high carbon footprint associated with fertilizer production and application, especially when growing corn as a feedstock, causes traditional bioprocesses to have a relatively high carbon footprint. To alleviate some of these challenges, researchers have proposed cyanobacteria and algae as alternative microorganisms to be used in bioprocessing, and have demonstrated photosynthetic production of fuels, plastics, and pharmaceuticals.^7^ However, these systems are still limited by slow growth rates and the relatively inefficient energy conversion of photosynthesis.^8^ To overcome these shortcomings, and with the expectation of cheaper and cleaner electricity in the intermediate future, various electromicrobial production (EMP) processes have been proposed and demonstrated (Fig. 1).

Although nomenclature for bioelectrochemical systems varies in the literature, we define EMP processes as any process that converts CO_2_ into a value-added product (*i.e.*, contains some form of primary production), uses electricity as the primary source of energy driving that transformation, and uses microbes to produce the final product. Perhaps most notable are systems based on Knallgas (aerobic hydrogen-oxidizing) bacteria, such as *Cupriavidus necator*, which uses molecular hydrogen (H_2_), produced by the electrolysis of water, to fix CO_2_. *C. necator* has historically been studied for production of its natively-produced polymer polyhydroxybutyrate (PHB)^9^ and of biomass for use as a single cell protein.^10^ More recently, *C. necator* has been engineered to produce other carbonaceous products including fuels and commodity chemicals.^11–13^ As an alternative, formatotrophic microorganisms have been employed, in which formic acid produced from the electrochemical reduction of CO_2_ is used as an energy source or assimilated by microbes to produce value-added products.^14–16^ Naturally formatotrophic microbes such as *C. necator* have been studied for this purpose,^17^ as have organisms engineered to express formate-assimilating pathways.^18^ Two-step systems have also been developed based on bio-acetate as an intermediary molecule, in which CO_2_ and H_2_ are consumed by the acetogen *Sporomusa ovata* to produce acetate, which is then converted by a heterotroph such as *E. coli* to produce various value-added products.^19^ Methanotrophs have also been proposed for the production of bioproducts including PHB.^20,21^ Methane and methanol, both potential feedstocks for methanotrophic bacteria, can be produced from CO_2_ through a variety of means using renewable electricity.^22–24^ In an attempt to obviate the need for an electrochemically-derived mediator molecule, electroautotrophic systems have been proposed in which carbon fixation is driven by direct electron transfer via reversible electron conduit proteins such as those found in *Shewanella oneidensis*.^25,26^

To date, research efforts have focused primarily on studying the fundamental metabolisms that permit EMP processes or on engineering metabolic pathways to enable production of specific products in relevant microbial chassis. Despite these successful bench-scale demonstrations, progress towards scaled and integrated processes has remained limited. Moreover, rigorous calculations of productivity and efficiency limits that can enable comparisons among EMP processes have been elusive, in part due to significantly different operating conditions across laboratories. Physics-based models that capture relevant phenomena (microbial growth and productions and consumption of species, acid/base reactions, gas/liquid mass transfer, *etc.*) can enable like-to-like comparisons across EMP processes. Additionally, such models are necessary to quantify design and operation strategies that optimize performance and to identify process parameters that limit productivity and efficiency.

However, EMP systems rely on subprocesses, such as electrocatalysis and carbon capture, that are outside the purview of most literature that focuses on the microbial and biochemical reaction engineering components of these processes. While metabolic efficiencies, productivities, and yields of these systems may be compared, these analyses do not consider differences in electrocatalytic efficiencies and productivities that affect the viability of the process as a whole. Hence, development of end-to-end process models that rely on the material and energy balances quantified in individual reactor models is necessary for a comprehensive analysis of the relative merits of EMP process options.

Life-cycle assessments (LCAs) are tools for quantifying the environmental impact of products and processes across their entire life cycle in relevant categories including greenhouse gas emissions, human and environmental health effects, and resource depletion. LCAs, which follow the standards set by ISO 14040 and 14044,^27,28^ aggregate and analyze material and energy flows as well as emissions from every step in the supply chain within a given system boundary and quantify the impact of a process in the desired impact categories. LCAs aid in decision-making in process design as they can be used to evaluate the environmental impacts of multiple alternatives and inform strategies to lower their footprints. Because EMP systems have been proposed as more sustainable alternatives to traditional bioprocesses, conducting LCAs on these systems is a crucial tool in assessing these claims.

Here, we present a detailed LCA of three major EMP process options relying respectively on acetate, H_2_, and formate/ic acid as mediator molecules and compare their impacts to a traditional bioprocessing scheme relying on corn-derived glucose (Fig. 1). We chose biomass, enzymes, and lactic acid as example products to represent the breadth of use-cases and production schemes for which EMP systems may be utilized. To enable our analysis, we developed two-phase bioreactor models that describe microbial growth and product formation, acid/base reactions, gas/liquid mass transfer, gas and liquid phase flow, and active pH control. The models are used to evaluate the effects of reactor parameters and operating conditions on critical performance metrics including productivity, titer, and material and energy efficiency, and are coupled to process models that present a complete picture of material and energy demands for the EMP processes.

Our results demonstrate that EMP systems can achieve a smaller carbon footprint than traditional bioprocessing strategies provided the electric grid is composed of >~90% renewable energy sources. In addition, EMP systems require significantly less land, which can help alleviate the “food vs. fuel” trade-off associated with crop-based bioprocesses. For each of the three products considered, the H_2_-mediated Knallgas bacteria system achieves the lowest overall global warming potential, indicating that this EMP strategy may be best-suited for industrial efforts with current technology. The tight coupling between our life cycle impact and reactor and process models revealed environmental hotspots that are key sites for future engineering and research efforts. Our analysis demonstrates the utility of integrating reactor, process, and life cycle impact models for comprehensively evaluating biotechnological processes. Together, the presented models, methodology, and analysis provide a framework for analyzing EMP systems that can help enable working, scalable, and sustainable electromicrobial production processes.

## Computational Methods

### System overview and governing equations

All bioreactor models assume well-mixed gas and liquid phases that are exchanged at fixed liquid- and gas-phase dilution rates. In the liquid phase, we consider, where relevant, dissolved CO_2_, dissolved H_2_, dissolved O_2_, bicarbonate anions (HCO_3_^−^), carbonate anions (CO_3_^2−^), protons (H^+^), hydroxide anions (OH^−^), sodium cations (Na^+^), chloride anions (Cl^−^), formic acid HCOOH), formate (HCOO^−^), acetic acid (H_3_C_2_O_2_H), acetate anions (H_3_C_2_O_2_^−^), lactic acid (H_5_C_3_O_3_H), lactate anions (H_5_C_3_O_3_^−^), enzyme (E), and microbes (X). In the gas phase, we consider CO_2_, H_2_, and O_2_. By neglecting ammonium/a species, we have assumed they are fed in excess to the liquid phase as NH_3_.

The well-mixed phases are assumed to have sufficient convective mixing such that no concentration gradients are formed. Such an open, well-mixed system must satisfy mass conservation, given generally for the liquid phase as

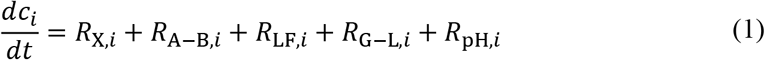

and for the gas phase as

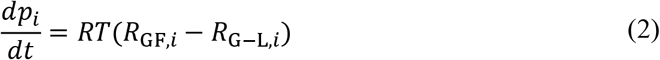

where *C*_*i*_ is the concentration, *p*_*i*_ is the partial pressure, *R*_*i*_ is the net volumetric rate of formation and consumption due to microbial growth (X), acid/base reactions (A–B), liquid or gas flow (LF/GF), gas/liquid mass transfer (G–L), and pH control (pH) for species *i*. The operating temperature is given by *T*, and *R* is the gas constant. Note that the gas phase species are assumed to follow ideal behavior and that the liquid and gas volumes in the reactor are equal.

### Microbial growth and product formation

Microbial growth occurs in the well-mixed liquid phase and is responsible for the production of more cells and the consumption or production of several chemical species. These reactions are compiled in *R*_X,*i*_. We assume that the kinetics of carbon fixation (or acetate uptake, in the case of acetotrophic growth) represent the upper bound on the biomass and product formation rates because all carbon-containing molecules produced by the cell are derived from the carbon-fixing metabolism. Hence, we assume that the combined rate of biomass and product (lactate or enzyme) formation (moles carbon per volume per time) is dependent on the molar biomass carbon concentration (*C*_X_) and the specific growth rate (*μ*). For lactate, this results in

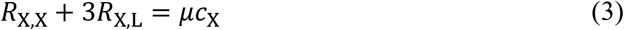

where the factor of 3 precedes *R*_X,*L*_ because lactate is a 3-carbon molecule. For the enzyme, the analogous equation is given by

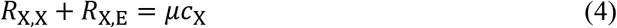

We define the fraction of carbon diverted to biomass as

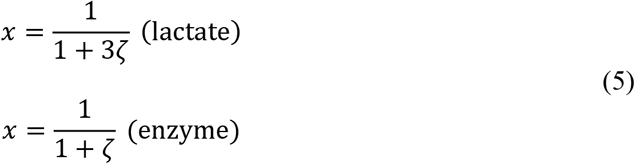

where *ζ* is the stoichiometric ratio of products to cells in, for example, the generic biomass equation given by

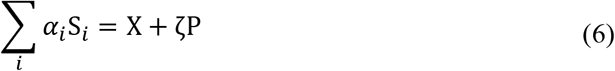

where S is a generic substrate and P is a generic product. We assume *x* is an engineerable parameter (*e.g.* by tuning the expression levels of different enzymes) and calculate *ζ* according to

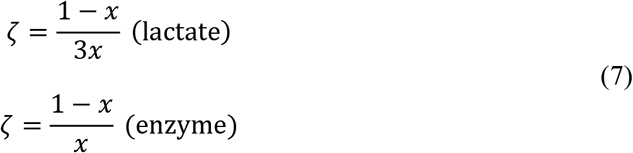

Hence, the biomass growth rate (*R*_X,X_) and product formation rate (*R*_X,L_, *R*_X,E_) are given by

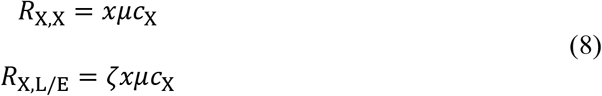

and consumption or production of other molecules (*e.g.* O_2_, H_2_, CO_2_, *etc*.) is written as

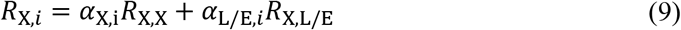

where *α*_*i*_ < 0 if the species is consumed in the reaction following standard convention.^29^

Microbial growth kinetics are defined using the Monod model with dependencies on each potentially growth-limiting substrate. The equations for aerobic formatotrophic (F), aerobic hydrogenotrophic (H_2_), anaerobic acetogenic (A), and aerobic acetotrophic growth (Ac) are given as

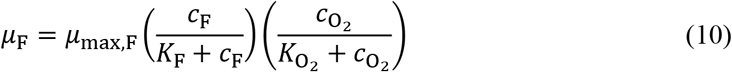

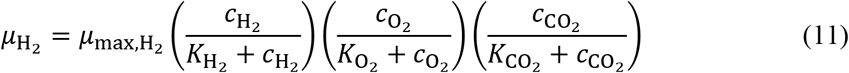

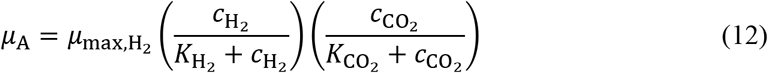

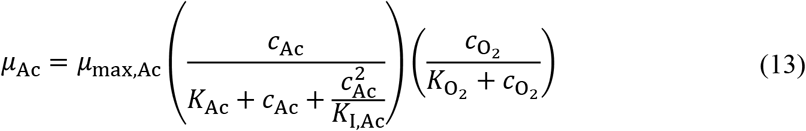

where *μ*_max_ is the maximum specific growth rate of the organism when all fixed carbon is diverted to biomass and *K*_*i*_ is the Monod constant for substrate *i*. Note that acetotrophic growth includes an Andrews/Haldane inhibition term (*K*_I,Ac_) to account for growth defects associated with high acetate concentrations reported previously.^30^

### Biomass and product yield

We use a combination of experimental values and stoichiometric and energetic calculations to determine the yields of biomass and products on different carbon and energy sources. In all cases we assume that enzyme yield 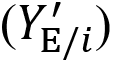 is equivalent to biomass yield 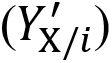 and that enzymes have approximately the same chemical composition as biomass.

#### Formatotrophic (aerobic) growth

For formatotrophic growth with O_2_ as the terminal electron acceptor and formate as the energy and carbon source (note that formate is completely oxidized and CO_2_ is fixed via the Calvin cycle in *C. necator*), the biomass reaction is written as

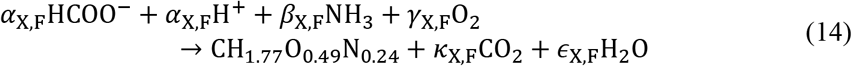

where CH_1.77_O_0.49_N_0.24_ represents cell mass (molar mass ~25 g mol^−1^). From stoichiometry,

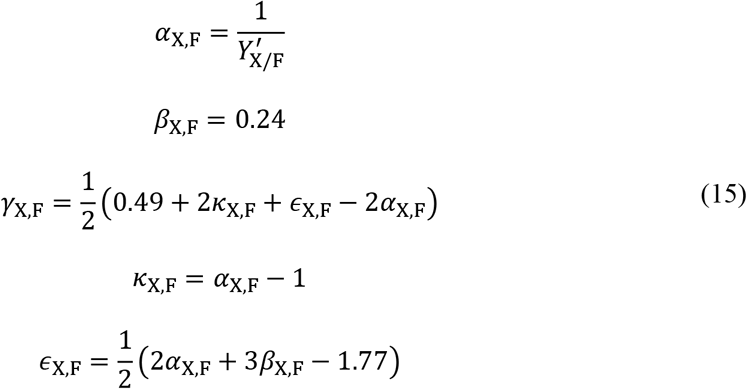

where 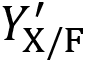 is the molar yield of biomass on formate, which we define according to a previously described empirical relationship,^16,17^

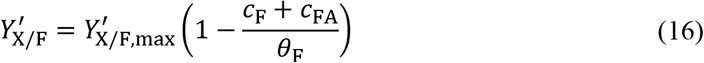

where *θ*_F_ is a fitting parameter that represents the maximum formate/ic acid concentration at which cells can grow.

The lactate formation reaction is written as

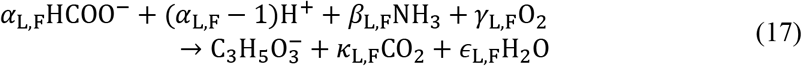

Relying on stoichiometry,

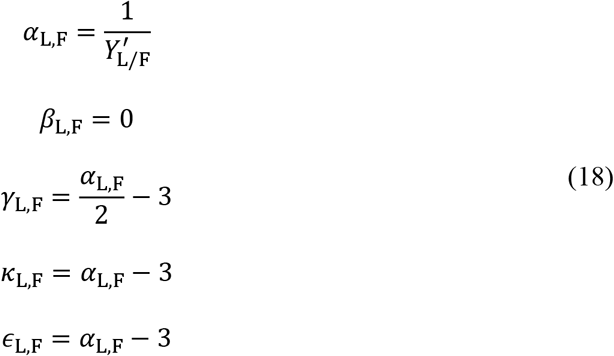

where 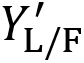 is the molar yield of lactate on formate. To determine this value, we follow the stoichiometry and energetics of carbon fixation via the Calvin cycle to lactate as follows. Microbes support energy carrier (NADH and ATP) regeneration by using NAD^+^-dependent formate dehydrogenases to catalyze the reaction

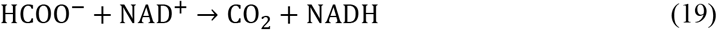

NADH is then used to regenerate ATP following aerobic respiration (oxidative phosphorylation):

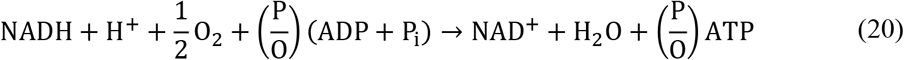

where P/O is the oxidative phosphorylation ratio (typically 2–3). When using the Calvin cycle to fix CO_2_, seven ATP and five NADH are consumed to fix three CO_2_ molecules into one pyruvate molecule:

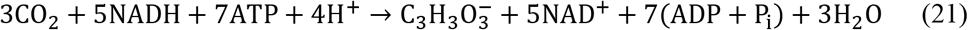

Pyruvate is then converted to lactate via lactate dehydrogenase according to

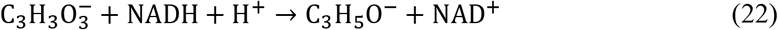

The resulting overall reaction for lactate production (using a P/O ratio of 2.5) is given by

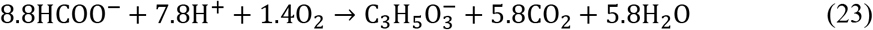

Hence, the maximum theoretical molar yield of lactate on formate is ~0.11 mol mol^−1^. Because the molar cell yield 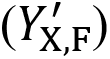 is influenced by the formate concentration due to a variety of toxicity effects in *C. necator*, we include this dependency for lactate as well:

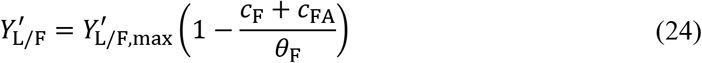

#### Hydrogenotrophic aerobic (Knallgas) growth

We use the same formulation as that for formatotrophy to describe biomass growth and product formation, but we modify the stoichiometry to account for the different energy source. The biomass equation is written as

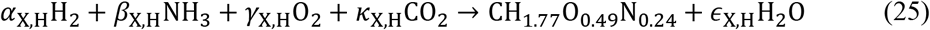

resulting in the stoichiometric relationships given by

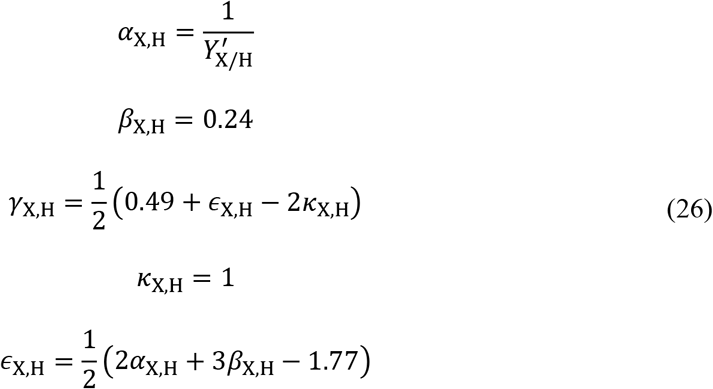

The lactic acid production reaction is written as

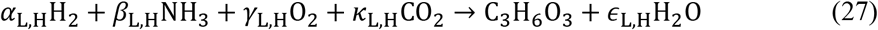

with stoichiometry given by

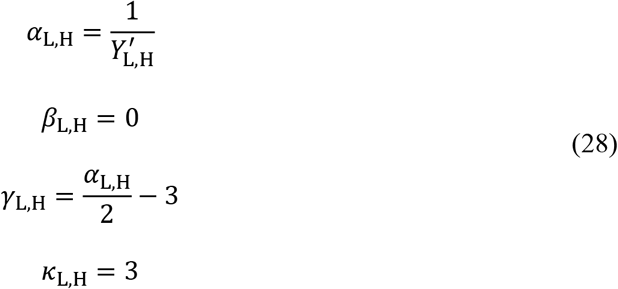

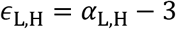

We determine the lactic acid production yield on 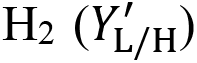 following the same method as for formate, resulting in:

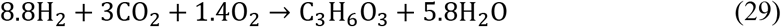

The equivalent theoretical yield of lactate on H_2_ and formate is because H_2_ and formate oxidation both result in the reduction of one molecule of NAD^+^ to NADH.

#### Acetogenic (anaerobic) growth

Anaerobic growth of acetogens relies on conservation of energy from the production of acetic acid,

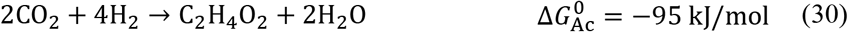

to drive anaerobic biomass formation given by

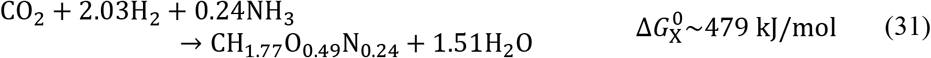

Following McCarty,^31^ we determine the ratio of acetic acid to biomass production (*ζ*_Ac_) using

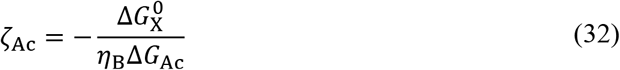

where *η*_B_ is the biological energy transfer efficiency (~0.6 across diverse microbial species). We calculate Δ*G*_Ac_ according to

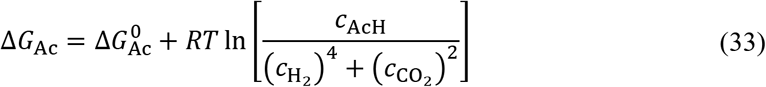

to account for deviation from standard-state conditions. Hence, the overall biomass equation for anaerobic acetogenic growth is written as

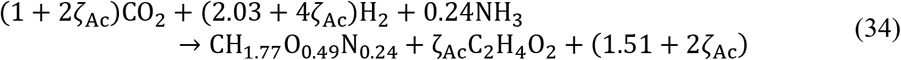

Here, the adjustable parameter *x* is unnecessary because reported growth rates already account for acetate production as it is an intrinsic part of the acetogenic metabolic process and the ratio of biomass to acetate is fixed by thermodynamics.

#### Acetotrophic (aerobic) growth

The biomass equation is written as

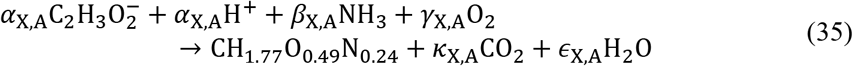

with stoichiometry given by

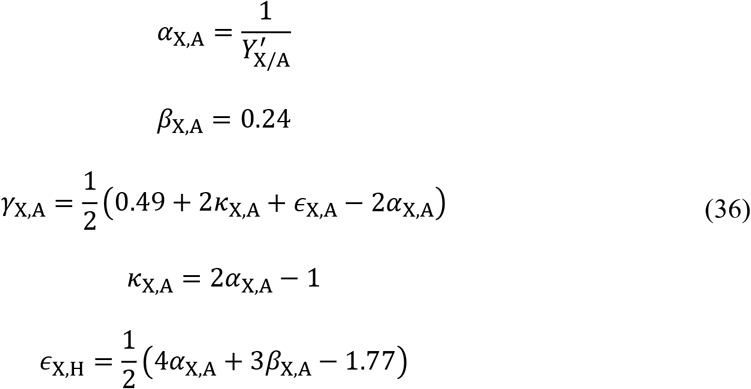

The lactate-forming reaction is written similarly,

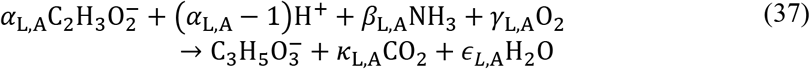

resulting in stoichiometry given by:

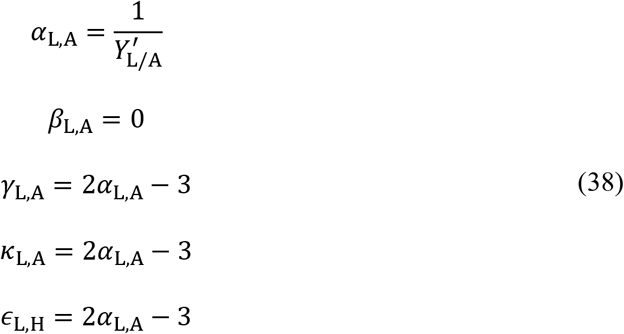

To determine the yield of lactate on acetate 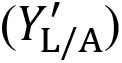, we follow the stoichiometry and energetics of acetate assimilation and oxidation through the glyoxylate shunt. Acetate is first activated to acetyl-CoA according to

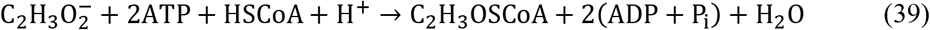

Note that here we’ve combined equations for ATP hydrolysis due to acetyl-CoA synthetase (resulting in AMP) and due to recombination with AMP resulting in 2 ADP. Acetyl-CoA is passed through the glyoxylate shunt to produce oxaloacetate and regenerate energy carriers, resulting in the net reaction given by

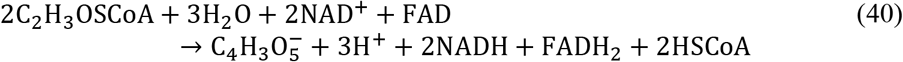

We assume oxaloacetate is converted to lactate via phosphoenolpyruvate and pyruvate with the net reaction

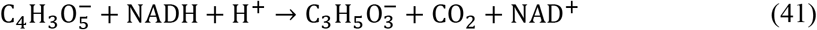

Using the P/O ratio of 2.5 for NADH (as above) and 1.5 for FADH_2_, the resulting net reaction for acetate conversion to lactate is given as

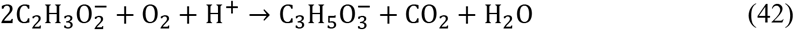

Hence, we use a theoretical molar yield of lactate on acetate 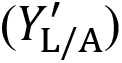 of 0.5 mol mol^−1^.

### Growth rate dependence on pH

We use a previously described model to describe the effects of pH on microbial growth:^16,32^

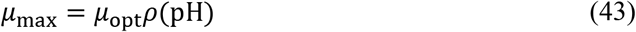

where *μ*_opt_ is the growth rate at optimal conditions and *ρ*(pH) are written as

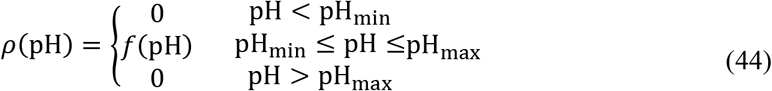

Here, *p*H_min/max_ is the range of pH over which microbial growth is observed, and the function *f*(pH) is

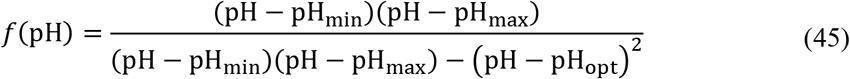

where pH_opt_ is the optimal pH for growth.

### Acid/base reactions

The acid/base bicarbonate/carbonate, formic acid/formate, acetic acid/acetate, lactic acid/lactate, and water dissociation reactions shown below occur in the liquid phase and are treated as kinetic expressions without assuming equilibrium:

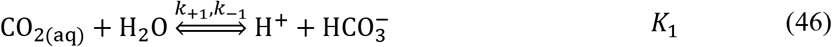

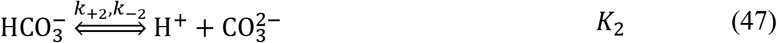

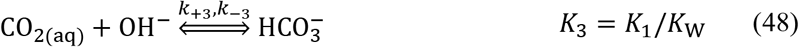

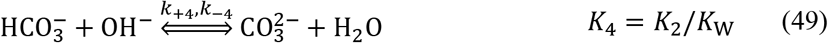

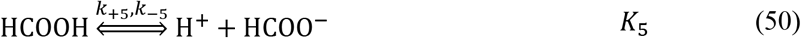

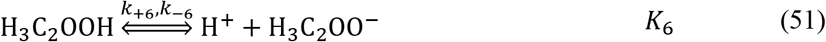

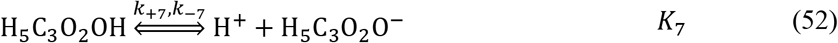

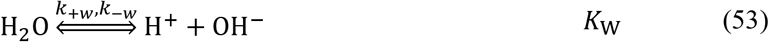

where *k*_+*n*_ and *k*_−*n*_ are the forward and reverse rate constants, respectively, and *K*_*n*_ is the equilibrium constant for the *n*th reaction, given by

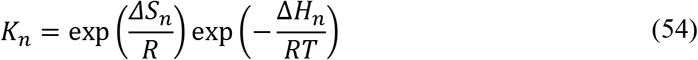

Source and sink terms resulting from these reactions are compiled in *R*_A–B,*i*_, written as:

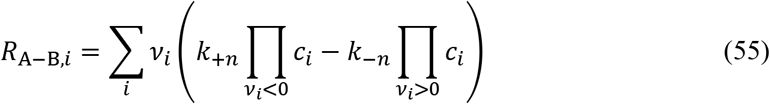

where *v*_*i*_ is the stoichiometric coefficient of species *i* for the *n*th reaction and reverse rate constants (*k*_−*n*_) are calculated from

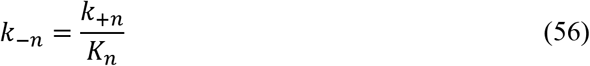

### Liquid and gas flow

Liquid media is fed to and extracted from the well-mixed liquid phase at a constant dilution rate, resulting in a feed term written as

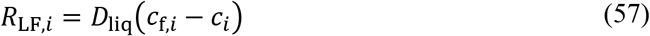

where *D*_liq_ is the liquid dilution rate (defined as the inverse space time, or volumetric flow rate divided by reactor volume). We assume the feed stream is free of microbes. We similarly define a feed term for the gas phase according to

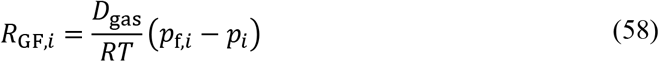

where *D*_gas_ is the gas dilution rate.

### Gas-liquid mass transfer

Gas fed to the reactor results in mass transfer to the liquid phase according to

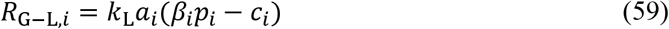

where *k*_L_*a*_*i*_ is the volumetric mass-transfer coefficient on the liquid side of the gas/liquid interface, and *β*_*i*_ is the Bunsen solubility coefficient. Volumetric gas/liquid mass transfer coefficients can be calculated from first principles^29^ or by using correlations that depend on the system geometry. For O_2_, we use the correlation developed by Vasconcelos *et al.* for stirred tank reactors with a height that is twice the diameter,

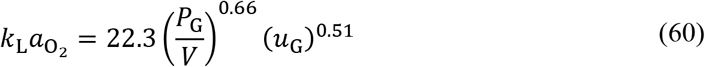

where *P*_G_/*V* is the specific power input (in units W m^−3^) and *u*_G_ is the superficial gas velocity (in units m s^−1^). We relate *u*_G_ to the gas phase dilution rate using

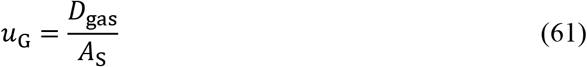

where *A*_S_ is the specific surface area of sparging holes in the reactor (units m^−1^). In our model, we assume a value of 5.6 m^−1^ to make a gas dilution rate of 100 hr^−1^ correspond to a superficial gas velocity of 0.05 m s^−1^, and we use the correlation above to determine the power demand necessary to achieve a given gas/liquid mass transfer rate.

To calculate the *k*_L_*a* value for CO_2_ and H_2_, we use

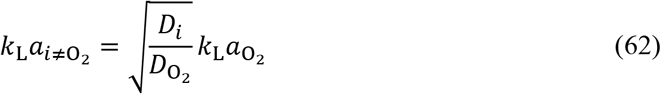

where *D*_*i*_ is the diffusivity of species *i* following Meraz *et al.* to account for differences in the mass transfer coefficient (*k*_L_).^33^

We calculate the equilibrium solubility of CO_2_, O_2_, and H_2_ according to the empirical relationship for the Bunsen solubility coefficient (*β*),

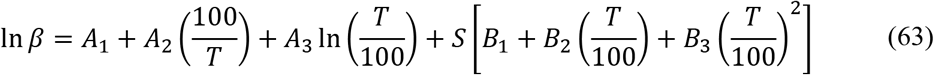

where *A*_*n*_ and *B*_*n*_ are fitting parameters and *S* is the medium salinity (in units g kg^−1^ water).

### pH control

A feedback control loop is included in the reactor to maintain an optimal pH for microbial growth by adding 1 M hydrochloric acid or 1 M sodium hydroxide solutions where appropriate. The manipulated flow rate variable (units hr^−1^) is defined as

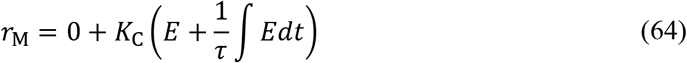

where *K*_C_ is the controller gain, *E* is the error, and *τ* is the controller reset time. The error (*E*) is defined according to

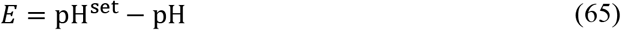

where pH^set^ is equivalent to pH_opt_. The resulting pH control flow is given by

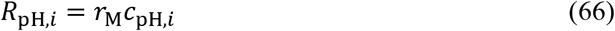

where c_pH,*i*_ is 1 M for H^+^/Cl^−^ (acid addition) or 1 M for OH^−^/Na^+^ (base addition).

### Reactor model analysis

We defined a normalized dilution rate (*δ*) according to

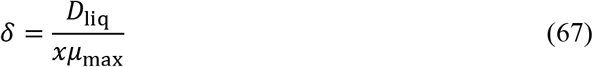

to account for the fact that the maximum growth rate is reduced by diversion of carbon to the product. The reactor productivity can then be calculated as

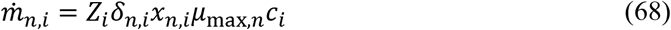

where *Z*_*i*_ is the molar mass of product *i* and the subscript *n* refers to a particular process. For the acetogenic system, we calculated the full-system productivity by accounting for flow through both reactors using

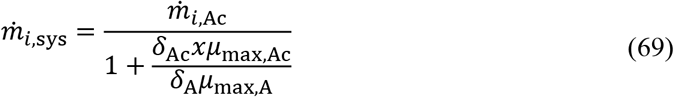

where the subscripts “Ac” and “A” refer to the acetotrophic and acetogenic reactors, respectively.

We calculated the energy efficiency of each reactor system according to

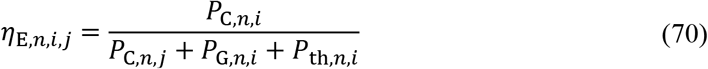

where the subscript *j* refers to the substrate, *P*_C_ is power embodied in the formation of species *i* or *j*, *P*_G_ is the power demand from mixing and gas/liquid mass transfer, and *P*_th_ is the power necessary to heat the liquid feed from room temperature to the operating temperature. We define the power of formation of a chemical species as the Gibbs free energy change per volume per time associated with the complete combustion of the chemical species following Claassens *et al.*^34^:

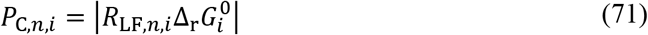

for liquid-phase species and

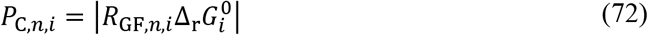

for gas-phase species. We note that these formulations mean that we have assumed residual substrate can be perfectly recycled and therefore represent upper bounds on the efficiency of the systems. In the formatotroph system, we include in the efficiency calculation the power necessary to concentrate formic acid effluent from the CO_2_ electrolyzer to the feed concentration of the reactor, which we define as

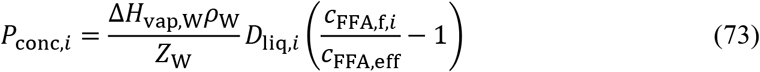

where Δ*H*_vap,W_, *ρ*_W_ and *Z*_W_ are the heat of vaporization, density, and molar mass of water, respectively, *C*_FFA,f,*i*_ is the total concentration of formate and formic acid in the feed stream for the system producing product *i*, and *C*_FFA,eff_ is the total concentration of formate and formic acid in the effluent of the CO_2_ electrolyzer. Overall efficiency for the acetogen-mediated process is calculated simply by taking the product of the efficiencies of each of the two individual reactors.

When determining the energy demand of full processes (in units kWh/kg product), we calculated the power demand for substrate formation as

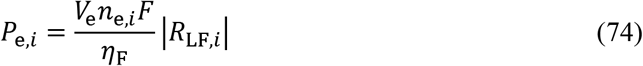

for the liquid phase substrates (formate/ic acid), and as

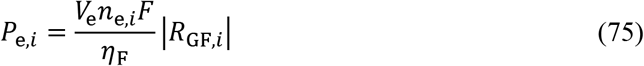

for gas phase substrates (H_2_). In each equation, *V*_e_ is the operating voltage of the electrolyzer producing species *i* (H_2_ or formate/ic acid), *n*_e_ is the stoichiometric ratio of electrons to electrolyzer product, *F* is Faraday’s constant, and *η*_F_ is the Faradaic efficiency of the electrolyzer.

### Reactor model implementation

All equations are solved using the MUMPS general solver in COMSOL Multiphysics 5.4. Model parameters are listed in Table S1.

### Life Cycle Analysis Goal and Scope Definition

This life cycle assessment was carried out according to the standards in ISO 14044.^27^ The open source life cycle assessment software openLCA version 1.10.3 (https://www.openlca.org/)^35^ was used to aggregate life cycle inventory data and apply impact assessment methods. The Product Environmental Footprints Dataset^36^ was used to obtain most background life cycle inventories while others were aggregated from literature as needed. Unless otherwise stated, the analysis was made indifferent to the exact location of the process. MATLAB was used to develop an impact model sensitive to changes of various variables and parameters studied.

The primary goal of this LCA is to predict the performance of three electromicrobial production systems (labelled as the Knallgas bacteria-based system, the formatotrophic system, and the acetogenic system) with regard to two sustainability metrics: global warming potential and land occupation. The LCA compares these systems to each other as well as to a traditional bioprocess using corn-derived glucose as a feedstock for a heterotrophic bacterium. A secondary goal of this analysis is to determine the specific limitations, bottlenecks, and environmental hotspots of each proposed EMP system. The final goal of this analysis is to integrate the life cycle impact model with the bioreactor models developed to create a tool enabling the eco-design of EMP processes.

### Functional Unit and System Boundaries

The production of four products is considered: biomass, industrial enzymes, lactic acid, and polylactic acid. The life cycle impact analysis ends at the production of each product in unprocessed form. For most products, downstream processing is not considered, as the processing of a given product would be identical for each system studied. For the production of biomass, the functional unit is 1 kg biomass. For industrial enzyme production, the functional unit is 1 kg of enzyme unpurified from the cell pellet. For lactic acid, the functional unit is 1 kg of lactic acid at a concentration of 100 g/L.^37^ For polylactic acid, the analysis ends at the production facility gate. Despite not considering end of life processing of the products, biogenic carbon is not considered as sequestered carbon, and all biogenic carbon is assumed to decompose to carbon dioxide.

### Process Modelling and Life Cycle Inventory

Material and energy requirements for the process are obtained from the results of the EMP reactor models and are sub-divided into the following categories: electricity generation for the EMP system; carbon dioxide direct air capture; ammonia production; other required nutrients and pH control agents; electrolyzer materials; and plant and bioreactor construction. In addition, a corn-derived glucose-fed *E. coli* process is modelled, in which glucose production is added as a process category. Carbon dioxide flows are explicitly considered in the EMP models. For all other nutrient requirements, the medium is assumed to be recycled such that 95% of input materials are consumed by the bioreactor. We assume a C:N:P ratio of 50:10:1 and base calcium, magnesium, and sulfur requirements on the elemental composition of *E. coli*.^38^

We assume each major process in the system draws electricity from a grid composed of coal, natural gas, hydropower, nuclear, photovoltaic, and wind derived energy. The composition of the grid is treated as a variable in our impact assessment model. The life cycle inventories of these six electricity sources are obtained from the Product Environmental Footprints (PEF) dataset. Direct air capture of carbon dioxide via temperature-vacuum swing adsorption is modelled based on Duetz and Bardow’s analysis of industrial-scale plants operated by Climeworks.^39^ The carbon capture process uses an amine-on-silica adsorbent and is powered by electricity obtained from the grid. Two possible routes for ammonia synthesis are considered, both involving the Haber-Bosch process. In one route, hydrogen for ammonia synthesis is obtained from steam methane reforming (SMR). In an alternative route, hydrogen is obtained from electrolysis of water drawing electricity from the grid (green ammonia). Material and energy requirements and life cycle impacts of ammonia production by both methods are adapted from Singh *et al*.^40^ A mix of ammonium phosphate (from phosphoric acid) and ammonium chloride (from hydrochloric acid) is supplied to the bioreactor to maintain the assumed C:N:P ratio. Life cycle inventories for phosphoric acid, magnesium sulfate, and calcium chloride are obtained from the PEF dataset. The pH is controlled in the bioreactor by addition of hydrochloric acid and sodium hydroxide, which are obtained through the Chlor-alkali process and rely on electricity from the grid. Energy requirements and life cycle impacts are derived from Garcia-Herrero *et al*.^41^ Electrolyzer material requirements are adapted from previous literature^42–44^ and the life cycle inventories associated with each component are obtained from the PEF database. The lifetime of the electrolyzers is assumed to be three years.

The process productivities obtained from the reactor models are used to determine the total bioreactor volumes required to produce the functional unit of a given product. Stainless steel bioreactors are used, with material requirements calculated based on the design of Mobius Bioreactors from EMD Millipore. The impacts of the bioreactor and the plant facility are due primarily to producing the required construction materials—stainless steel for the bioreactor and concrete and steel for the plant, assuming a constant amount of concrete and steel per square meter of facility area.^45^ The area of facility space required per aggregate volume of the bioreactors is based on the Natureworks lactic acid production facility in Blair, NE. Steel, stainless steel, and concrete life cycle inventories are all obtained from the PEF database. We assume a reactor lifetime of eight years and a plant lifetime of thirty years.

Glucose for the heterotrophic process is obtained from the hydrolysis of corn starch, and life cycle inventories of glucose production are obtained from the PEF dataset. Ammonia requirements for corn production are obtained from Ma *et al*.^46^ and the life cycle inventories for glucose production are adjusted to account for reduced carbon emissions in the case of green ammonia production.

### Life Cycle Impact Assessment

Global warming potentials were calculated according to the 2013 IPCC model for 100-year global warming potential and are expressed in kilograms of CO_2_-equivalents [kg CO_2_-e].^47^ The land use footprint is calculated using the ReCiPe (H) 2016 method, which weights the impact of various types of land use by their impact on biodiversity.^48^ The units of land use are expressed as m^2^.year crop equivalents, representing the weighted land use needed to produce a given functional unit of product per year.

### Sensitivity Analysis

All parameters used in the development of the bioreactor models and life cycle analysis (e.g., growth rates, reactor lifetimes, solar electricity GWP) other than physical properties (molecular weights, heat capacities, etc.) were independently altered by +/−30% and the global warming potential of each process was recalculated. The ratio of the global warming potential of each EMP process (formatotrophic, Knallgas, and acetogenic) and the global warming potential of the heterotrophic process in each scenario was taken to be the metric of interest to evaluate the sensitivity of each parameter. The parameters that caused the largest deviation of this ratio from the equivalent ratio for the base case value of all parameters were taken to be the most critical parameters in the study (a 10% deviation of this ratio from the base case value was used as a cutoff). A list of all parameters studied can be found in Supplementary Table 1.

## Results and Discussion

### Reactor models reveal trade-offs in productivity and efficiency across processes

Below we compare the productivity (Fig. 2a–c), titer (Fig. 2d–f), and efficiency (Fig. 2g–i) of the three EMP processes producing biomass, enzyme, and lactic acid, describing trends for each system, and making comparisons between base case conditions.

**Figure 2.**
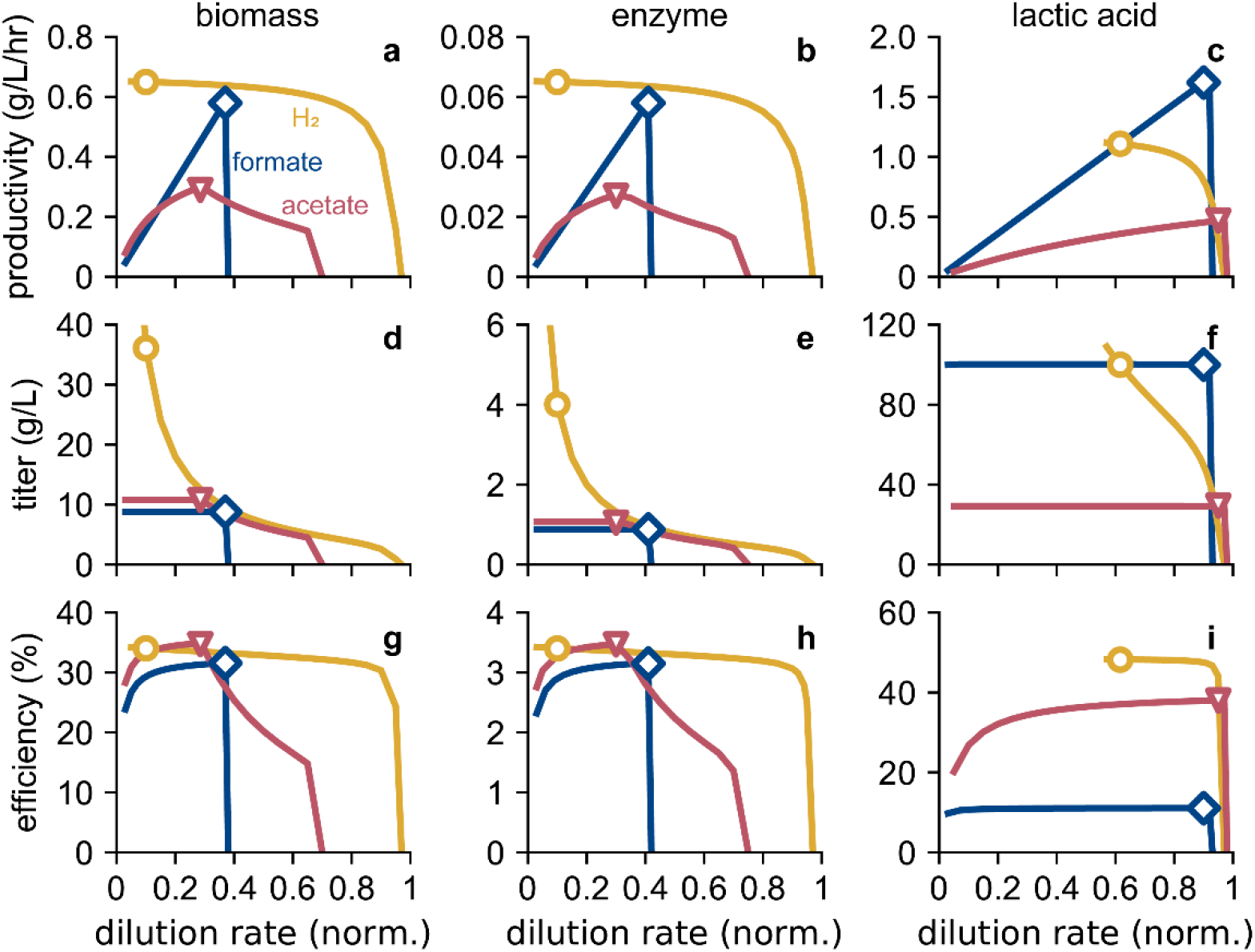
EMP reactor performance. Productivity (**a**, **b**, **c**), titer (**d**, **e**, **f**), and energy efficiency (**g**, **h**, **i**) as a function of normalized dilution rate (*δ*) for the three EMP systems producing biomass (**a**, **d**, **g**), enzyme (**b**, **e**, **h**), and lactic acid (**c**, **f**, **i**). Base case conditions are indicated by blue diamonds (formate-mediated), yellow circles (H_2_-mediated), and red triangles (acetate-mediated). The color scheme in all panels follows that in (**a**).

Because formate is fed in the liquid phase, the formatotrophic system follows the standard trend of initially increasing productivity for each product followed by a rapid decline as cell washout occurs (Fig. 2a–c). However, washout occurs well before the dilution rate exceeds the maximum growth rate; this is due to the limitation on productivity imposed by O_2_ gas/liquid mass transfer. As the dilution rate increases, the formate feed rate exceeds the consumption rate limit imposed by O_2_ mass transfer, causing toxic build-up of formate in the reactor and an associated reduction in the biomass growth rate, resulting in cell washout. Mass transfer limit-induced washout dynamics are also observed for biomass (Fig. 2a) and enzyme (Fig. 2b) production in the acetogenic system, although the system behavior for acetate is slightly different than for the formate case due to the different strategies for modeling acetate and formate toxicity. In the case of lactic acid production in the acetogenic system, the biomass growth rate, as opposed to O_2_ gas/liquid mass transfer, sets an upper bound on productivity, so neither acetate accumulation nor its impact is observed (Fig. 2c).

Productivity of biomass, enzyme, or lactic acid in the Knallgas bacteria-based system does not follow the typical trend because all substrates necessary for growth are fed via the gas phase. Hence, productivity is only slightly dependent on the liquid phase dilution rate until washout begins to occur at a normalized dilution rate of ~0.85 (Fig. 2a–c). Instead, for the Knallgas system, product titer is controlled by the liquid dilution rate, enabling a wide range of achievable product titers (Fig. 2d–f). We did not explore the full dilution rate range for this system in the case of lactic acid production because below a normalized dilution rate of ~0.55, the basic NaOH feed necessary to maintain a neutral pH results in a salinity >3%, which we defined as toxic to the cells. For both the formatotrophic and acetogenic systems, product titer is fixed by the feed concentration, as is standard for substrates supplied in the liquid phase (Fig. 2d–f). The lactic acid titer in the acetogenic system could not exceed ~30 g/L due to the same salinity-induced toxicity concerns as for the Knallgas system. In all product cases, for each EMP process, efficiency follows the same trend as productivity. Hence, for a given process option, maximizing productivity also maximizes efficiency.

We therefore selected base-case operating conditions by maximizing productivity for each system (Table 1). A minimum normalized dilution rate of 0.1 was arbitrarily set for the Knallgas system because it has a wide dilution rate range at which the productivity is roughly equal, and for lactic acid production in the formatotrophic system we assumed a concentrated (12 M) formate feed stream to achieve an industrially-relevant titer (100 g/L). Considering biomass first, the achievable productivity is highest for the Knallgas system at ~0.65 g/L/h, ~10% and ~50% higher than the productivities of the formatotrophic and acetogenic systems, respectively (Fig. 2a). The former difference is due to the ~13% higher biomass yield on O_2_ with H_2_ as the energy substrate than with formate and the fact that the H_2_ gas/liquid mass transfer limit is slightly lower than the O_2_-imposed limit. The acetogenic system, in contrast, is primarily limited by the acetate production rate of the acetogen, which grows ~4-fold slower than Knallgas and formatotrophic bacteria.

**Table 1.**
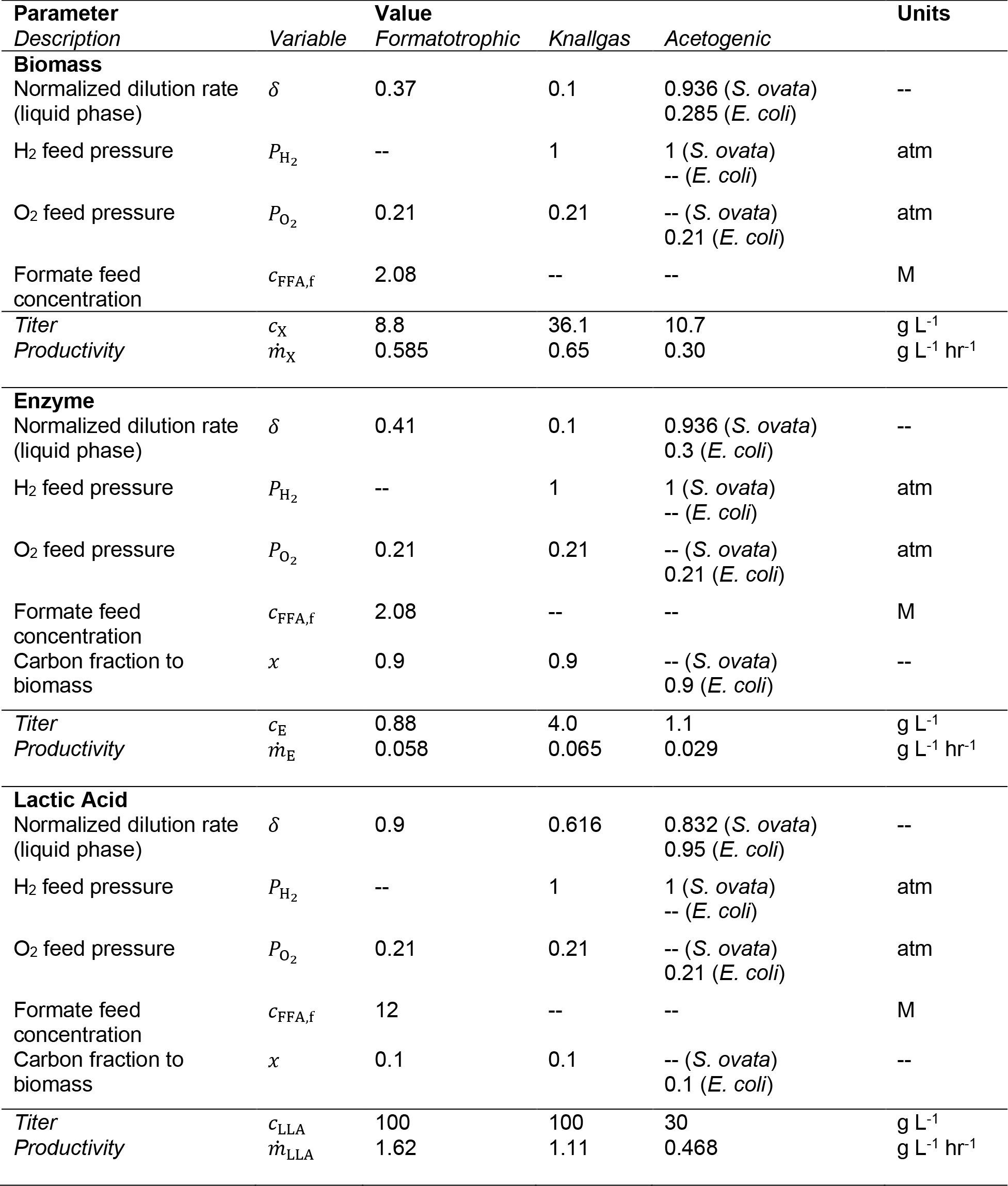
Base case operating conditions.

The Knallgas system also achieves the highest biomass titer (~36 g/L vs. ~10 g/L for each other process) because the titer is fully controllable by the liquid-phase dilution rate for this system (Fig. 2d). These trends are also true for the enzyme production case, although the productivity, titer, and efficiency are all ~10-fold lower than for biomass because we assume only 10% of the fixed carbon is diverted to enzyme production (Fig. 2b, e). For both biomass and enzyme formation, the efficiency of each EMP process is remarkably similar (~32-35% for biomass production, Fig. 2g) and are dominated by the metabolic efficiency, defined as the ratio of energy embodied in the product to energy embodied in the main substrate. That these efficiencies are nearly equal, and that the acetate-mediated system maintains a slight advantage, is surprising given the remarkably different metabolic strategies for biomass (or enzyme) production.

In contrast to the case for biomass and enzyme formation, lactic acid productivity is highest with the formatotrophic system at ~1.6 g/L/h, ~46% and ~240% higher than the Knallgas and acetogenic systems, respectively (Fig. 2c). The Knallgas system is limited by the H_2_ gas/liquid mass transfer rate, while the acetogenic system remains limited by the acetogen growth rate. This enhanced productivity, however, comes with a significant energy penalty associated with concentrating the formate/ic acid feed stream. This results in a significantly lower energy efficiency for the formatotrophic system at ~11%, ~4-fold lower than the Knallgas system and ~3.5-fold lower than the acetogenic system. We note, however, that because the lactic acid titer in the acetogenic system is only ~30 g/L, additional energy will be required in downstream purification steps that are not considered in this efficiency calculation.

Several initial conclusions can be drawn from this analysis. First, both the Knallgas and formatotrophic systems can achieve higher productivities and efficiencies than the acetogenic system. The acetogen-based system does maintain advantages not captured in this analysis, including that a wider range of industrial microorganisms (*e.g. E. coli*, *Bacillus licheniformis*, and oleaginous yeasts) grow naturally on acetate, but bioengineering efforts could obviate this advantage in the future. Second, the solubility advantage of formate as a growth substrate compared to H_2_ is only relevant in cases where the O_2_ gas/liquid mass transport is a less stringent limit on productivity than H_2_ transport. This depends both on the ratio of H_2_ to O_2_ in the gas phase and the ratio of H_2_ to O_2_ consumed per unit of product. Under the conditions we explored here, the O_2_ mass transport limit is rate-determining for biomass production such that the Knallgas system achieves higher biomass productivity than the formatotrophic system (Fig. 2a). However, H_2_ mass transport limits lactic acid productivity for the Knallgas system, so the increased solubility of formate enables a higher lactic acid productivity (Fig. 2c). Although concentrating formate from the effluent of a CO_2_ electrolyzer represents a significant energy penalty, reducing efficiency (Fig. 2i), improvements in CO_2_ electrolysis reactor operation may overcome this challenge, as we discuss later. We also note that the gas mixture we have assumed for the Knallgas system is flammable. A nonflammable gas mixture would either require significantly less air (~0.24 atm as opposed to the assumed 1 atm), reducing productivity by decreasing O_2_ solubility, or significantly more H_2_ (~4.15 atm vs. the assumed 1 atm), increasing safety concerns associated with pressurized gases and likely increasing reactor and control systems complexity. These results indicate trade-offs in productivity, titer, and efficiency such that reactor models alone cannot identify a clearly-best EMP strategy. Moreover, upstream processes including energy-substrate generation (via either water or CO_2_ electrolysis), CO_2_ capture, ammonia production, NaOH and HCl production for pH control, and other considerations, require explicit attention as important drivers of material and energy demand for EMP processes. We therefore developed a complete process model (diagrammed in Fig. 3) for the EMP processes to understand material and energy flows for the full system, which we discuss next.

**Figure 3.**
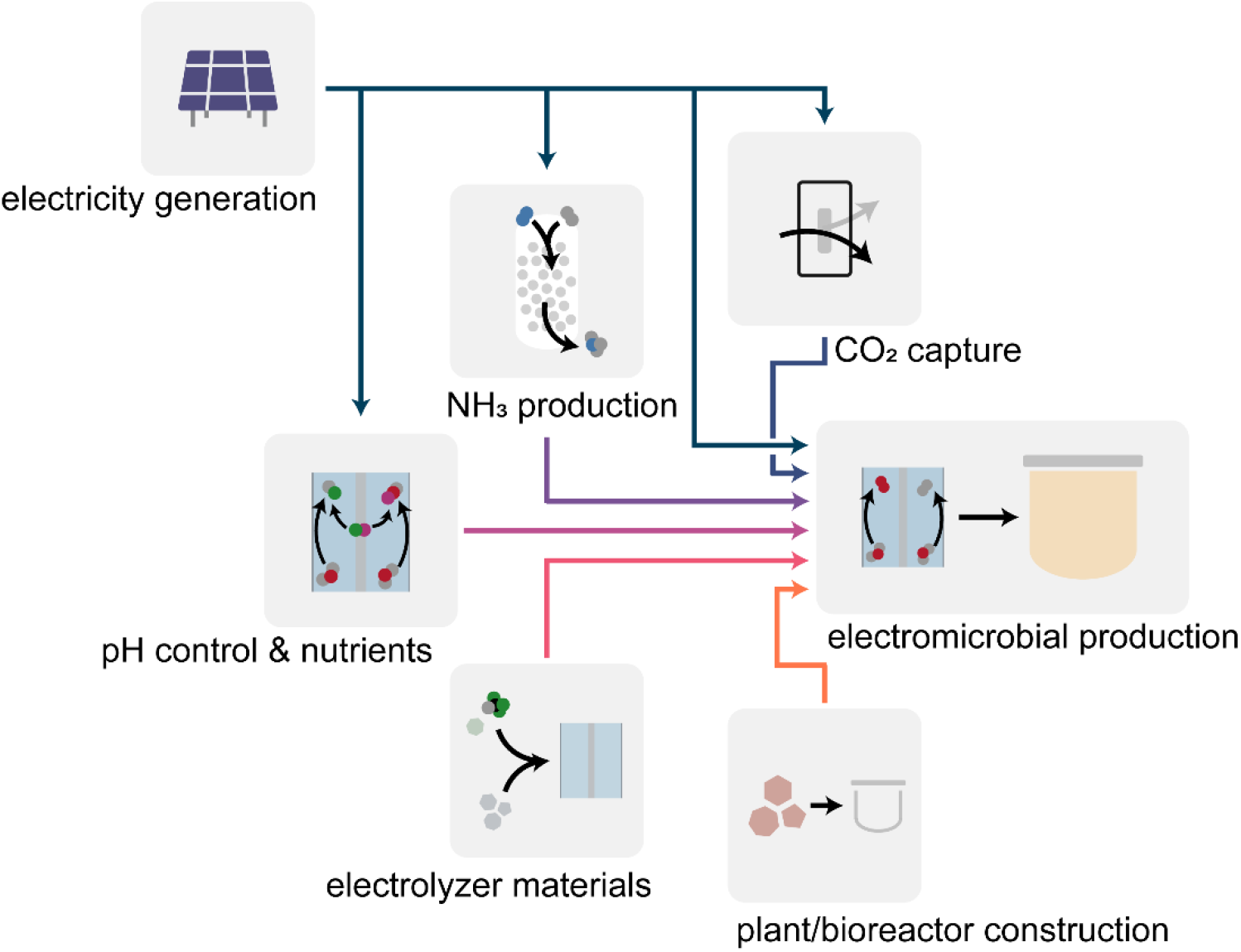
Schematic representation of the EMP system. Grid electricity (midnight blue) supplies electricity to the EMP reactors and supporting processes, including direct air capture of CO_2_ (blue), ammonia production via the Haber-Bosch process (purple), and the chlor-alkali process producing pH control agents (magenta). Mining and production of electrolyzer materials (pink) and materials for reactor and plant construction (orange) are also considered within the impact model.

### Energy requirements for EMP Processes

Considering the entirety of an EMP process there are five major energy demands: electrosynthesis of mediator molecules (H_2_ or formic acid), bioreactor energy demands (heating, gas-liquid mass transfer, etc.), direct air capture of carbon dioxide, green ammonia production, and production of NaOH and HCl for pH control through electrolysis of NaCl and water. The material and energy flows for each of the three systems are summarized in Table 2.

**Table 2.** Energy and material demand for biomass production.

| Process Demand | Value |  |  | Units |
| --- | --- | --- | --- | --- |
|  | *Formatotrophic* | *Knallgas* | *Acetogenic* |  |
| **total process electricity** | **53.8** | **28.4** | **31.5** | **kWh/kg CDW** |
| electrolysis & bioreactor electricity | 48.0 | 23.2 | 21.5 | kWh/kg CDW |
| CO<sub>2</sub> | 1.93 | 1.76 | 1.98 | kg CO<sub>2</sub>/kg CDW |
| NH<sub>3</sub> | 0.179 | 0.179 | 0.214 | kg NH<sub>3</sub>/kg CDW |
| total NaOH and HCl | 0.15 | 0 | 3.23 | kg/kg CDW |

In all cases, electrolysis to produce H_2_ or formate contributes the majority of the energy demand of each system over the entire process. Even though the acetogenic EMP system has the lowest energy requirements when considering only electrolysis and bioreactor operation (as seen in Table 2), the Knallgas bacteria system has the overall lowest energy demand of the three systems when considering the entire electromicrobial production process. This is caused by a combination of lower CO_2_ and NH_3_ consumption, and the lack of required pH control in the Knallgas bacteria system. The increased carbon and nitrogen requirements of the acetogenic process stem from the “wasteful” production of *S. ovata* biomass.

The Knallgas bacteria system requires no pH control because conversion of H_2_ and CO_2_ to biomass involves no net consumption or generation of protons. The formatotrophic system requires only a relatively small amount of NaOH to balance the pH due to the formic acid feed. Our model predicts the acetogenic system requires substantial pH control, as conversion of CO_2_ into acetate lowers the pH of the *S. ovata* medium (requiring addition of basic solution) while conversion of acetate to biomass raises the pH of the heterotroph reactor (requiring addition of acidic solution). Owing to the substantial amount of electricity required by the chlor-alkali process to produce NaOH and HCl, pH control accounts for 12.4% of the total electricity required by the acetogenic EMP process.

### Global Warming Potential

The global warming impacts of all components shown in Fig. 3 were calculated as outlined in the methods section for each of the three EMP systems and the traditional glucose-fed process. For the case of a wind-powered process, the global warming potential broken down by process categories is shown in Fig. 4. It should be noted that other means of clean electricity production (such as thin-film photovoltaics and hydropower) have roughly equivalent life cycle emissions per kWh produced, and therefore would lead to similar results. To study general trends regarding the potential of each process alternative, 1 kg of biomass is chosen as the product and functional unit as a baseline comparison.

**Figure 4.**
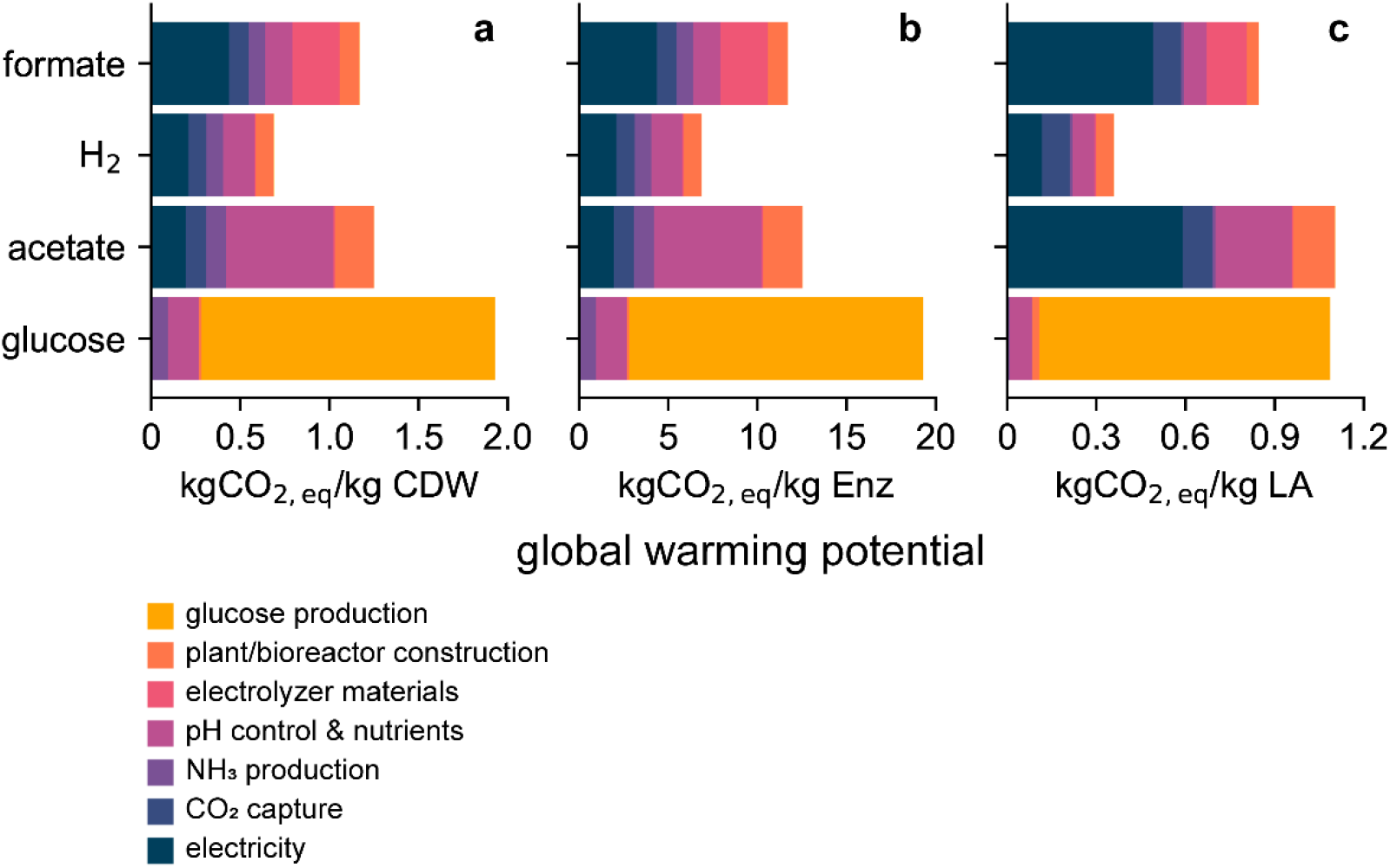
Global warming potential of EMP and traditional bioprocesses. Global warming potential for the three EMP systems and traditional heterotrophic system for the production of (**a**) biomass, (**b**) enzymes, and (**c**) lactic acid, broken down by process category.

Our impact model shows that all three proposed EMP systems have the potential to have a lower global warming potential than that of the corn-based glucose-fed bioprocess, given a clean electricity source. Our analysis indicates the Knallgas bacteria system has a lower overall global warming potential (0.69 kg CO_2_-eq./kg biomass) than both the formatotrophic system and acetogenic system (1.17 and 1.25 kg CO_2_-eq./kg biomass respectively). The reduction in greenhouse gas emissions associated with the electromicrobial production system compared to the heterotrophic system stems from the low emissions of the individual components of the EMP systems when drawing energy from a low-impact energy grid. The high-impact agricultural production of corn and other crops as feedstocks in bioprocesses contributes the largest share of the global warming potential of these systems. While the carbon emissions associated with fertilizer production can be reduced with increased clean energy (as calculated in our impact model), the large amount of nitrous oxide emissions due to fertilizer application will not be affected by this change, leading to a relatively large global warming potential of traditional bioprocesses. Therefore, in a clean-electricity dominated scenario, the Knallgas bacteria-based EMP system will have a GWP 64% lower than a glucose-fed process (Fig. 4).

Although both rely on the same microorganism in our model (*C. necator*), the formate-mediated electromicrobial system will have a larger global warming potential than a hydrogen-mediated system. CO_2_ electrolysis to formate occurs at lower current densities (140 mA/cm^2^ vs. 1 A/cm^2^) and with higher overpotentials (>2 V vs. ~0.8 V) compared to water electrolysis, resulting in an increased carbon footprint due to an increased demand for electrolyzer materials (*e.g.* Ir, Pt, Nafion, *etc.*) and increased energy consumption. We further describe the effects of potential improvements to this system in the later section titled “Engineering Targets for Formate Electrolysis”.

The greatest environmental hotspot of the acetogen-based system compared to the others is due to the production of NaOH and HCl for pH control. The chlor-alkali process that produces NaOH and HCl is an energy-intensive electrolytic process, and therefore contributes a substantial carbon footprint. Even when running the chlor-alkali process with clean electricity, NaCl production and other processing steps still contribute to the carbon footprint of pH control.^41^ However, there are a couple options to help alleviate this constraint. For example, engineering the acetogen-based process to take place in a single reactor could address the problem of pH control because the combined biochemical reactions result in no net generation or consumption of protons. The key impediment to this solution is the strict oxygen sensitivity of acetogens such as *S. ovata*^49^ and the requirement for oxygen in assimilation of acetate as a sole carbon source.^50^ However, for certain applications, this may be achievable. *S. ovata* has recently been evolved to tolerate low concentrations of oxygen.^51^ If paired with a heterotroph producing a product traditionally produced by fermentation such as butanol,^52^ microaerobic conditions would be suitable to achieve high yields. Therefore, the aeration conditions of the two organisms could be similar enough to warrant their co-culture in a single reactor.

Further transitions to a clean energy grid will likely reduce the carbon footprint of EMP processes due to a combination of effects too granular to be captured in our model. The life cycle carbon footprint of solar energy production, for example, will likely fall as silicon production and purification processes begin to use cleaner energy. Emissions due to transportation along the supply chain will likely fall due to increased use of electric vehicles. As such changes continue to occur, it is in principle feasible for electromicrobial production processes to achieve full carbon neutrality. The carbon footprint of glucose-based bioprocesses, however, is unlikely to achieve full carbon neutrality. Cleaner methods of fertilizer production and electrified processes for farming machinery and glucose processing will indeed lower the carbon footprint of conventional bioprocesses. However, the primary source of greenhouse gas emissions in corn production is due to the application of fertilizers, as nitrogenous fertilizers are partially degraded to nitrous oxide, a greenhouse gas with 298-fold higher global warming potential than of CO_2_.^46,47^ Further transitions to a clean electric grid and electrified processing, then, are more likely to decrease the global warming potential of EMP processes than of heterotroph-based processes.

We extended our life cycle impact analysis to the other two products modeled, industrial enzymes and lactic acid. In the case of an industrial enzyme as the product of interest, the trends largely follow that of biomass. Assuming the industrial enzyme product is intracellular, effects of titer do not impact the energy demand as low-energy separation methods (*e.g.*, settling, filtering, centrifuging) are possible. Therefore, the similar trends for GWP in enzyme production and biomass production, scaled due to the relative yields of each, are expected.

In the case of lactic acid production, the trends between EMP systems are similar to those of biomass production, with Knallgas bacteria-based production of lactic acid exhibiting the lowest global warming potential of the systems studied. In this system, we define the functional unit to be a solution of lactic acid at a concentration of 100 g/L. The formatotrophic system does have a relatively higher GWP than the Knallgas system; CO_2_ electrolysis achieves only a limited titer of formate/ic acid (~2.08 M in our base case) because parasitic oxidation of formate can occur at the anode. Hence, concentrating the effluent is necessary to achieve a high productivity and desired titer of lactic acid resulting in an additional energy penalty associated with the distillation process. Similarly, the acetogen-based system produces lactic acid at a low titer according to our bioreactor models, requiring energy-intensive downstream purifications to achieve the desired 100 g/L lactic acid titer. Our impact model suggests that the acetogen-based system will have an overall higher GWP than a traditional glucose-based system due to these effects. However, we do assume an energy-intensive evaporation process is required for lactic acid concentration. Lower-energy methods of separation, such as extraction, may produce lower global warming potentials. This, however, is beyond the scope of our study. We have also calculated a cradle-to-grave life cycle global warming potential of polylactic acid (PLA) made from lactic acid in each of these processes (Supplementary Fig. 1), assuming the PLA is composted at the end-of-life. We have found that PLA made from EMP-generated lactic acid will have lower life-cycle greenhouse gas emissions compared to petroleum-based plastics such as polystyrene (PS) and polyethylene terephthalate (PET), assuming a sufficiently high yield (see Supplementary Note 1).

Importantly, the data shown in Fig. 4 assume 90% of the fixed carbon is converted to lactic acid, which matches the yield commonly achieved by lactic acid fermentation from glucose ^53^. This high yield of lactic acid, achievable due to the high yield of fermentation products during anaerobic growth, may not be achievable in EMP systems. All three EMP systems considered (based on hydrogen-oxidizing, formatotrophic, or acetotrophic metabolism) require respiration, suggesting the high yield of lactic acid may not be achievable. We expand on the effect product yield has on the viability of EMP systems in the following section.

### Effect of Electric Grid on Global Warming Potential of EMP Systems

The dominant factor affecting the environmental sustainability of electromicrobial production systems is the source of energy due to the high electricity demand of each system. We therefore studied how the electricity grid composition affects the global warming potential associated with each system (Fig 5a). Although our impact model can input an electric grid composition comprising several sources (coal, natural gas, hydropower, nuclear, photovoltaic, and wind), to simplify the results we assumed an electric grid comprised of some combination of wind power and natural gas, defining the fraction of electricity derived from wind power as the “percent of grid renewable.” It should be noted that the present electric grid composition of the United States (as of 2019) has a global warming potential of about 450 g CO_2_/kWh, roughly equivalent to a grid that is based 100% on natural gas.^54^ As shown in Fig. 5a, even the Knallgas bacteria system, which has the lowest GWP of the three schemes, will not have a lower GWP than a traditional glucose-based system unless over 90% of the electric grid comes from renewable resources. These “breakeven” values are 95% and 97% renewable electricity for the acetogenic and formatotrophic system, respectively.

**Figure 5.**
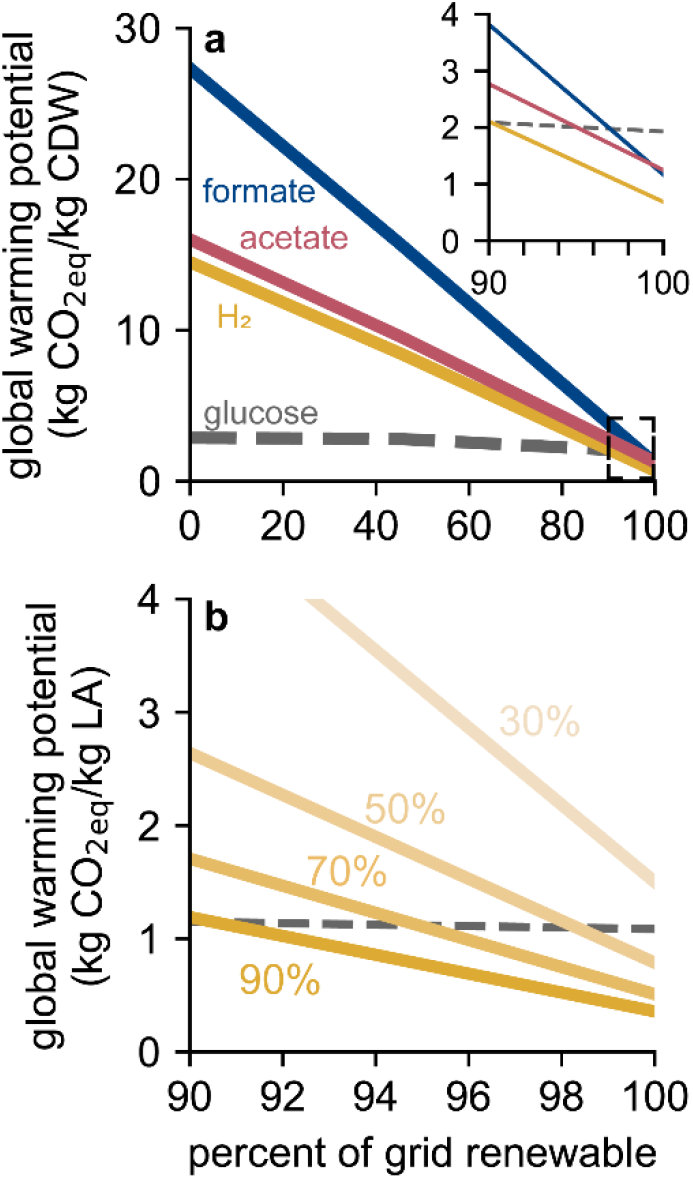
Effect of electricity grid on global warming potential. (**a**) Global warming potential for the production of biomass for the formate-(blue), acetate-(red), H_2−_(Knallgas, yellow), and glucose-fed (traditional bioprocessing, dashed gray) systems drawing electricity from a grid composed of variable fractions of wind power (renewable) and natural gas (non-renewable). Inset shows the >90% renewables region bounded by the dashed box. (**b**) Global warming potential for the production of lactic acid in the H_2−_fed (Knallgas) system (yellow) as a function of electricity grid compositions for variable carbon efficiencies (fraction of fixed carbon diverted to lactic acid) as well as global warming potential for the traditional glucose fermentation of lactic acid (dashed gray).

**Figure 6.**
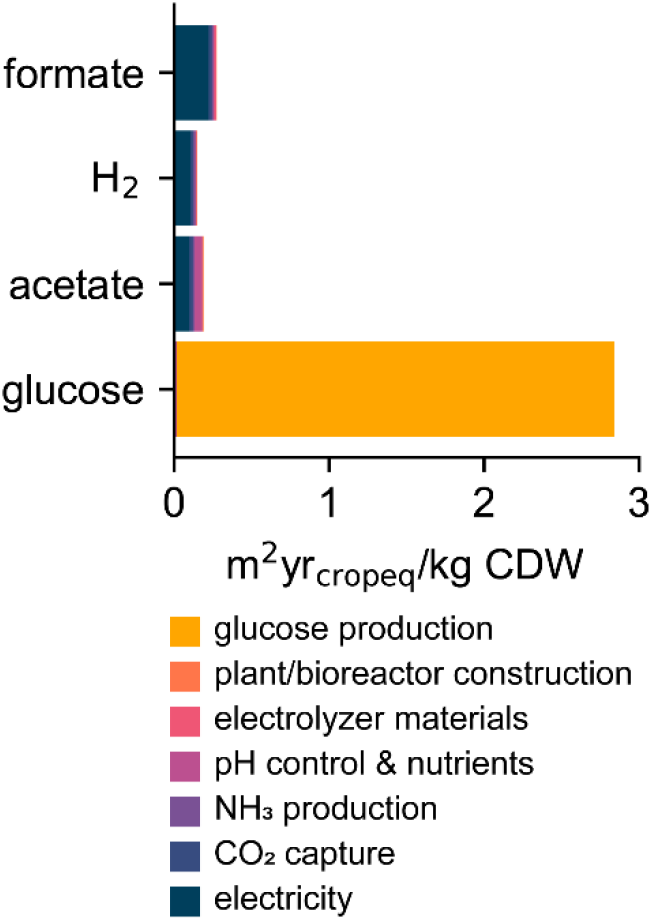
Land occupation footprints of EMP and traditional bioprocesses. Land occupation footprint as calculated by ReCiPe 2016 (H) midpoint method for the three EMP systems and traditional heterotrophic system for the production of biomass, broken down by process category.

Fig. 5b shows similar breakeven curves for the production of lactic acid in a Knallgas bacteria-based system at a range of carbon efficiencies (defined as the fraction of fed or fixed carbon diverted to lactic acid). As noted previously, lactic acid production in EMP systems has not yet been demonstrated, although other similar products (i.e. traditionally made through fermentation) such as isopropanol and butanol have been produced with moderate carbon efficiency.^19,55^ The carbon efficiency of product formation will play a large role in its global warming potential. At 90% carbon efficiency, lactic acid production in a hydrogen-fed system will attain environmental viability if at least 90% of the electric grid is composed of renewable resources. At lower carbon efficiencies, stricter requirements of the grid are necessary to achieve a lower GWP than traditional glucose-fed processes. At 30%, production of lactic acid through EMP will result in higher greenhouse gas emissions than the glucose-fed process (Fig. 5b) regardless of the electricity source. This result highlights the importance of maximizing the product yield as electromicrobial systems are developed and establishes a target yield of at least 50% of the theoretical maximum.

### Land Use

Our impact assessment demonstrates a significantly lower land use of EMP systems compared to traditional bioprocesses, even when using solar energy, the electricity generation method with the largest land occupation footprint. This result is expected as EMP processes have been proposed in part to alleviate the “food vs. fuel debate” that stems from the high agricultural land use of traditional bioprocesses and biofuels. The dominant factor determining the land occupation of EMP systems is the land used by solar panels required for electricity production, including electricity production for electrolysis, ammonia production, and HCl and NaOH production. Therefore, the land occupation impacts follow the same trends of total electricity use in Table 2. The weighted land occupation footprint of the Knallgas bacteria system is 0.15 m^2^·yr crop-eq./kg compared to 2.84 m^2^·yr crop-eq./kg, representing a 95% reduction in land use. The improved energetic efficiencies of lithoautrotrophic carbon fixation compared to photosynthetic carbon fixation is the primary driver of this large disparity.

We report a weighted land occupation footprint as described by the ReCiPe 2016 midpoint method, which weights different types of land use according to their impact on the environment. Solar panels may be deployed in many environments, including sparsely vegetated or urban land, and therefore will have lower land use impacts than agricultural land use. When discounting weighting factors, the raw land use of the Knallgas bacteria-based system is 0.25 m^2^·yr./kg biomass, representing an 11-fold decrease in land use compared to a glucose-fed system. These results are consistent with the energetic efficiencies of solar-to-biomass efficiencies of 9.7% for a *Cupriavidus necator* hydrogen-fed system and ~1% for photosynthetic plants reported by Liu *et al.*^55^ Although land use data from databases are generally recognized to be less reliable than greenhouse gas emission data, the land occupation footprint of both EMP systems and of the traditional bioprocess are dominated by the land requirements of solar panels and corn farmland respectively, both of which have well-studied data. Additionally, the calculated land requirements of EMP processes are over an order of magnitude smaller than that of heterotroph-based processes. We are therefore confident in our assessment than EMP processes will have a substantially lower land occupation footprint than current biomanufacturing methods.

### Intrinsically safer operation of the H_2_-mediated Knallgas system

Despite the lower GWP associated with the H_2_-mediated system, the flammable gas mixture fed to the reactor under base case operating conditions (1 atm H_2_ and 0.21 atm O_2_) may pose a significant barrier to adoption of this EMP strategy. We therefore evaluated intrinsically safer operation (ISO) of the H_2_-mediated system by adjusting the H_2_:O_2_ ratio in the gas phase such that the gas mixture was inherently non-flammable (defined as comprising an H_2_:O_2_ ratio of >19.76:1).^56^ Under these conditions, O_2_ gas/liquid mass transfer limits the productivity for each product, biomass, enzymes, and lactic acid. However, reactor productivities equivalent to that of the base case scenario can be achieved simply by increasing the total gas pressure while maintaining the inherently non-flammable gas ratio, so the GWP of the H_2_-mediated EMP process is not negatively impacted by ensuring intrinsically safer operating conditions. For biomass, enzymes, and lactic acid, the partial pressure of H_2_ must be 4.14 atm, 4.14 atm, and ~2.45 atm, respectively, to match the GWP of the base case scenario, pressures that are readily achievable with existing water electrolysis and bioreactor technology.

### LCA as an ecodesign tool: engineering targets for formate electrolysis

The formate-mediated EMP system is associated with a significantly higher GWP than the H_2_-mediated system due primarily to differences in electrolyzer performance. To identify engineering targets that must be met by CO_2_ electrolysis systems, we calculated the GWP of biomass and lactic acid production as a function of electrolyzer parameters (current density, *j*; energy efficiency, *η*; formate titer, *C*_FFA_), and we compared these results to the H_2_-mediated system operated under intrinsically safe conditions (Fig. 7).

**Figure 7.**
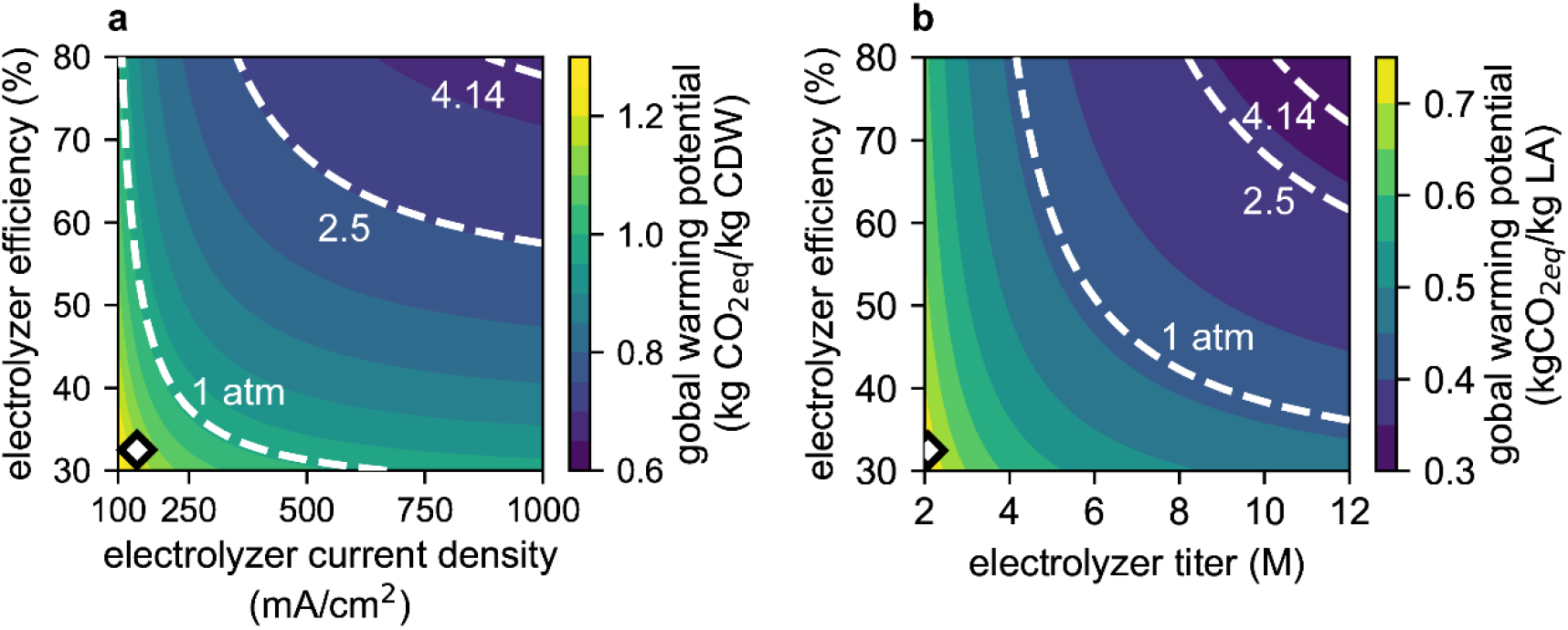
Effects of CO_2_ electrolysis operating parameters. Global warming potential of (**a**) biomass production with the formate-mediated system as a function of electrolyzer current density and electrolyzer efficiency and (**b**) lactic acid production with the formate-mediated system as a function of electrolyzer formate titer and electrolyzer efficiency. Overlaid white dashed lines in (**a**) and (**b**) correspond to the global warming potential of the intrinsically-safer H_2_-mediated system (as described in the text) operating at different H_2_ partial pressures. Point in (**a**) highlighted by the diamond (white fill, black outline) denotes the base case CO_2_ electrolysis operation using current technology. Point in (**b**) highlighted by the diamond (white fill, black outline) denotes the base case titer and efficiency of the CO_2_ electrolyzer, but (**b**) assumes an operating current density of 1 A/cm^2^.

Considering biomass first, base-case electrolysis operation (*j* = 140 mA/cm^2^, *η* = 32.5%) results in a significantly higher GWP than the H_2_-mediated system (Fig 7a). A current density of >~250 mA/cm^2^ and *η* >~40% is necessary to outcompete the H_2_-mediated system operating at an H_2_ partial pressure of 1 atm, while a current density in excess of ~900 mA/cm^2^ with *η* >75% is necessary to reach parity with an H_2_-mediated system operating at 4.14 atm of H_2_ (Fig. 7a). For lactic acid, in addition to operating at 1 A/cm^2^, the formate electrolyzer must achieve an efficiency of >~75% and a formate titer >~11 M to match the GWP of the H_2_-mediated system operating at 4.14 atm (Fig. 7b). Despite significant progress towards improving CO_2_ electrolysis performance in the past decade,^57^ these metrics represent extremely challenging targets that may be infeasible. Hence, H_2_-mediated EMP systems based on Knallgas bacteria appear to be better-suited for industrial adoption.

### LCA as an Ecodesign tool: effect of electric grid and reactor lifetime

Our approach of integrating a physics-based bioreactor model with a life cycle impact model can also provide a decision-making tool in designing electromicrobial processes at scale. Alone, the bioreactor model may provide reasonable estimates for productivities, titers, and energy efficiencies that can be achieved in scaled-up electromicrobial production processes, thereby providing a valuable tool in the design of such processes. However, these values alone provide little insight into real-world implications of these systems, particularly in terms of their environmental sustainability. Our approach of integrating a bioreactor model with a life cycle assessment framework provides the ability to contextualize the tradeoffs that may occur between efficiency and productivity and provide a single metric (*i.e*., global warming potential) by which to evaluate the sustainability of a particular EMP design. We highlight this utility by returning to the example of comparing a formatotrophic EMP system (high productivity, low efficiency) with that of a Knallgas-based system operating under inherently safer conditions and atmospheric pressure (low productivity, high efficiency).

All other impact model parameters held constant, the impact model can be reduced to:

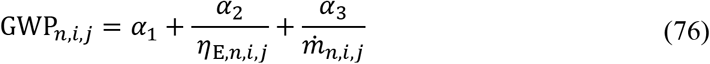

The parameters *α*_1_, *α*_2_, and *α*_3_ are obtained from the life cycle impact model and are dependent on myriad subordinate parameters. These parameters provide a weighting system for comparing the independent effects of energy efficiency and productivity. However, it is important to note that these parameters are not universally-set parameters but depend on several specifics of the process design.

For example, *α*_2_ is strongly dependent on the global warming potential of the electric grid from which energy is supplied (which in turn is dependent on the composition of that grid), while *α*_3_ is mostly dependent on specification of the reactor and plant infrastructure required for the EMP process, and therefore varies with parameters such as the reactor lifetime. Therefore, there is no universal answer for how to weigh the tradeoffs between efficiency and productivity, and therefore no universal solution for comparing the formatotrophic and the (inherently safer) Knallgas EMP systems. We can, however, provide a tool that considers the parameter landscape of these systems and shows under what conditions efficiency or productivity become dominant factors in minimizing the global warming potential of a system.

Fig. 8 demonstrates the effect of reactor lifetime and electricity global warming potential (which is in turn dependent on the electricity source) on relative greenhouse gas savings of either the Knallgas system or formatotrophic system. In our base case scenario, assuming a GWP of 9 g/kWh (the GWP associated with wind power) and a reactor lifetime of 8 years, the Knallgas system is preferred (yellow region in Fig. 8). This benefit is driven by the higher energy efficiency of this system. However, if the reactor lifetime is reduced, the weighting factor associated with the productivity of the system increases, as the reactor size per functional unit of product increases. Therefore, if the reactor lifetime becomes sufficiently short, the productivity of the system will become a more important factor in determining the GWP of the system, which will then favor the formatotrophic system (blue-shaded region in Fig. 8). Likewise, if the electricity global warming potential is decreased further, the formate-mediated system will be favored as the energy savings of the Knallgas system will become less important. In addition to demonstrating under which conditions either a H_2_- or formate-mediated system is superior in terms of minimizing greenhouse gas emissions, Fig. 8 demonstrates the novel utility of an integrated bioreactor/life cycle model in the eco-design of electromicrobial production systems.

**Figure 8.**
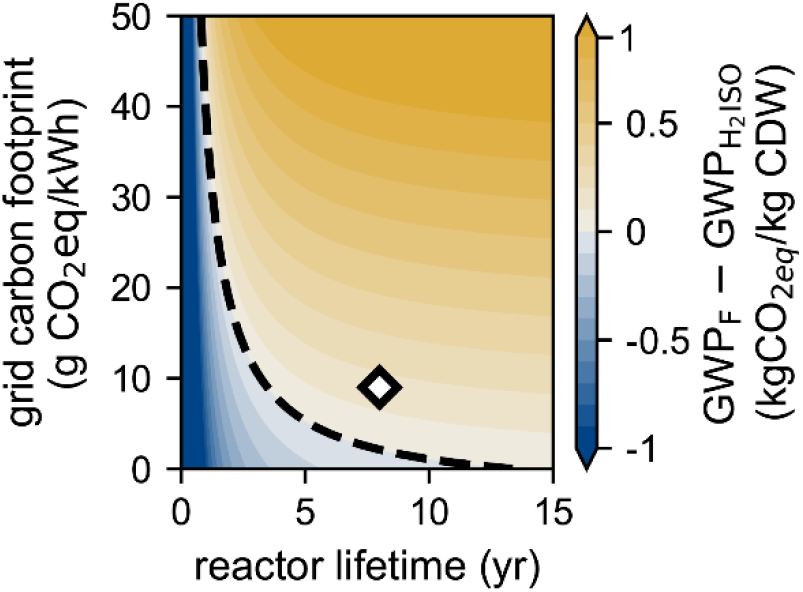
Life cycle impact-based break-even analysis. Difference in global warming potential of biomass production between formate-mediated and intrinsically safer H_2_-mediated EMP systems as a function of bioreactor lifetime and the electric grid carbon footprint. Base case conditions are highlighted by the diamond (white fill, black outline), and the break-even condition (*i.e.* no difference between the two systems) is denoted by the black dashed line.

### Assessment limitations and parameter sensitivity analysis

To investigate the impact of uncertainty on our model and conclusions, we performed a sensitivity analysis on the 77 individual parameters in our model (Table S1 in the supplementary information). We identified the most important parameters, defined as those for which a 30% change in the parameter value induced a >10% change in the global warming potential of the impacted process(es) for biomass production (Fig. 9). Of these, three (global warming potential of glucose, biomass yield on glucose, and global warming potential of wind-produced renewable energy) are outside the scope of reactor models and therefore do not affect the productivities or efficiencies of any of the electromicrobial production systems. These parameters do, however, impact the comparative life cycle assessment of these systems. Of these three parameters, the life cycle global warming potential of glucose production has the greatest potential to affect our assessment indicating the advantages of EMP systems over traditional systems. There is lack of consensus in the literature and databases on this value, which range between 0.75-1.2 kg CO_2_-eq./kg glucose, due to variations in corn growth methods, locations, and processing, as well as allocation methods.^58,59^ Therefore, the range shown in Fig. 9a does reasonably depict the uncertainty of our analysis. Therefore, if EMP systems are to replace traditional bioprocess methods, attention must be paid to the specifics of the heterotrophic feedstock production to ensure an accurate comparison. However, we do note that even in the “worst-case” scenario, all three systems do outperform a traditional bioprocess.

**Figure 9.**
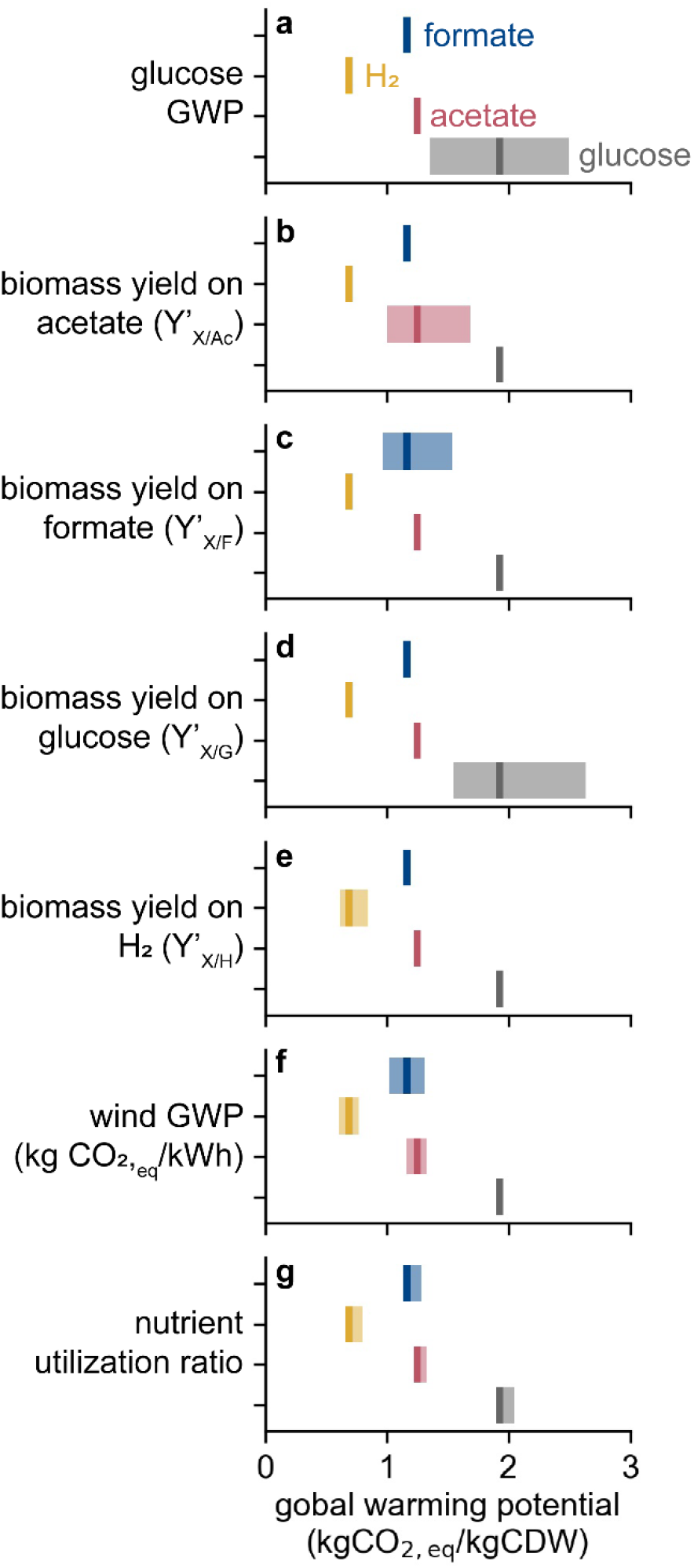
Parameter sensitivity analysis. Global warming potential dependence of producing biomass on +/−30% variation in (**a**) glucose production global warming potential, (**b**) biomass yield on acetate, (**c**) biomass yield on formate, (**d**) biomass yield on glucose, (**e**) biomass yield on H_2_, (**f**) wind-based energy production global warming potential, and (**g**) the nutrient utilization ratio for each process. Dark bars represent base case values, shaded bars represent range in global warming potential induced by variation.

The yield of biomass on glucose in the heterotrophic system can also significantly affect the analysis. However, although literature yields on glucose do vary slightly,^60,61^ a 30% deviation in this value is unlikely, reducing the uncertainty related to this parameter. The global warming production of electricity production (wind power in the base case) does impact the overall life cycle assessment, particularly in the formatotrophic system where the electricity demand is highest. However, due to the low carbon footprint of wind energy, even a 30% increase of this value would not significantly affect the viability of EMP systems in comparison to heterotrophic systems.

Biomass yields on acetate, formate, and H_2_ can all significantly impact the global warming potential of the relevant processes; however, the Knallgas bacteria system still outperforms the others even if the biomass yield on H_2_ is 30% lower than expected while the biomass yields on formate and acetate are 30% higher (Fig. 9). Notably, microbial growth rates do not significantly impact global warming potential mainly because gas/liquid mass transfer rates impose an upper bound on productivity (see discussion around Fig. 2). Because the electricity demand associated with achieving high kLa values is minimal compared to energy substrate generation via electrolysis (Table 1), productivity improvements via increased agitation or other strategies to enhance gas/liquid mass transfer rates are a straightforward strategy to reduce the carbon footprint of a given process. The final significantly impactful parameter is the nutrient utilization ratio, indicating that efforts to recycle unconsumed nutrients (especially ammonia) are also important for the viability of EMP (and traditional) bioprocesses.

The sensitivity analysis demonstrates that neither 30% variability in any single parameter nor any set of parameters is sufficient to dislodge Knallgas bacteria-based EMP systems as the process with the lowest global warming potential, although variation in some single parameters can result in re-ordering EMP processes: for example, a 30% higher yield on acetate enables the acetate-mediated system to outperform the formate-mediated system). However, all EMP processes outcompete glucose-based bioprocessing given 30% uncertainty in any single parameter, and concomitant variation in multiple parameters in particular directions is required for glucose-based systems to achieve parity with any of the EMP processes. This analysis indicates that our conclusions are robust to significant uncertainties in parameters used in our reactor, process, and life cycle impact models.

## Conclusions and outlook

In this study we have provided the first comparative life cycle assessment of electromicrobial production systems, an important step in their validation as promising bioprocessing methods. Our impact assessment of biomass production shows that all three EMP (formate-mediated, H_2_-mediated, and acetate-mediated) systems analyzed can lead to potential reductions in greenhouse gas emissions when compared to traditional heterotroph-based processes provided the electric grid consists of >~90% renewable energy sources. Furthermore, our analysis for enzyme and lactic acid production demonstrates the potential of a breadth of bioproducts while also elucidating important considerations (such as product yield) for the environmental viability of such systems. Through our analysis, we have shown that the H_2_-mediated, Knallgas bacteria-based system requires the least energy among EMP systems while achieving the lowest global warming potential. We also identified hotspots and potential improvements of both the acetogen- and formatotroph-based systems, demonstrating the utility of an integrated bioelectrochemical model/life cycle assessment framework in both analyzing and aiding the ecodesign of electromicrobial processes.

Our analysis focused on the engineering considerations of EMP systems, as well as their potential environmental impact. Detailed techno-economic evaluations are outside the scope of this present study. However, our model may provide a “first-pass” assessment of key economic considerations. For example, the key distinction between EMP processes and traditional bioprocesses are the differences in feedstock, with EMP processes utilizing electricity and carbon dioxide while traditional systems require sugars such as glucose. Assuming a future price of direct air capture of $100/tonne,^62^ we calculate the cost of feedstocks for the Knallgas EMP process will be lower than that of a glucose-fed process if the cost of electricity drops below $0.027/kWh. Although lower than the 2019 U.S.-average industrial electricity price of $0.067/kWh, the U.S. Department of Energy Sunshot 2030 initiative seeks to reduce the price of solar to $0.03/kWh by 2030.^63^ Furthermore, public policy affects these calculations. Current agricultural subsidies, for example, may impact the price of corn glucose as a feedstock,^64^ while proposed carbon pricing mechanisms such as a carbon tax would financially benefit EMP processes due to their potentially lower carbon footprints compared to traditional systems. Importantly, this basic assessment does not consider the cost of equipment and labor and cannot give a complete picture of the economic viability of EMP processes. Future techno-economic assessments should be performed to study these in detail. However, we can predict that in the coming decades, the price of electricity should not be a limiting factor in the cost of electromicrobial production.

Currently, electromicrobial systems are not environmentally viable due to the high carbon footprint of electricity generation. However, the climate crisis does necessitate a transition to a fully decarbonized grid in the coming decade. As of April 2021, it is the position of the present U.S. administration that the U.S. electricity grid will be composed of at least 80% renewable energy sources by 2030, with a fully decarbonized grid by 2035. However, while technically and economically feasible, meeting these ambitious yet necessary targets will require strong public policies, according to an analysis by Abhyankar *et al.*,^65^ including the passage of a Clean Electricity Standard (CES) by the U.S. Congress. Despite a large majority (63% of U.S. voters) supporting a Clean Electricity Standard, given the current makeup of the Congress passage of such legislation will likely require either the elimination of the Senate filibuster or passage through the budget reconciliation process.^66^ Our analysis therefore suggests that EMP systems can effectively reduce the negative environmental impacts of the biotechnology industry due its reliance on and competition with agriculture. However, policy concerns, rather than technological hurdles, are the primary obstacle towards the realization of these systems.

Electromicrobial production has the potential to “electrify” the biotechnology industry. However, our analysis suggests that electromicrobial production systems, if implemented in the United States today, would lead to higher greenhouse gas emissions compared to traditional bioprocesses due to the abundance of fossil energy sources in the current electric grid. Nonetheless, as the electricity grid continues to decarbonize in the coming decades, EMP will become an attractive alternative method of bioproduction. We suggest therefore that pilot-scale electromicrobial production systems of various value-added products be developed in the coming years, such that further scaling and distribution of these systems can be accomplished in the coming decades as the electricity grid becomes fully decarbonized.

## Author Contributions

**Conceptualization:** A.J.A., J.D.A.; **Data curation:** A.J.A., J.D.A.; **Formal analysis:** A.J.A., J.D.A.; **Funding acquisition:** D.S.C.; **Methodology:** A.J.A., J.D.A.; **Software:** A.J.A., J.D.A.; **Supervision:** D.S.C.; **Visualization:** A.J.A., J.D.A.; **Writing - original draft:** A.J.A., J.D.A.; **Writing - review & editing:** A.J.A., J.D.A., D.S.C.

## Acknowledgements

This work was supported by the Center for the Utilization of Biological Engineering in Space (CUBES, https://cubes.space/), a NASA Space Technology Research Institute (grant number NNX17AJ31G). A.J.A. is supported by an NSF Graduate Research Fellowship under grant number DGE 1752814. We thank Dr. Jacob Hilzinger (UC Berkeley) for feedback on the analysis, Marisa Watanabe for advice on schematics, and Dr. Paul Tol (Netherlands Institute for Space Research, SRON) for a helpful reference on accessible color schemes (https://personal.sron.nl/~pault/). The authors acknowledge this research was performed on unceded land of the Chochenyo-speaking Ohlone people.

## Notes

### Competing Interest Statement

The authors have declared no competing interest.

