## Supplementary Information for "A comparative life cycle analysis of electromicrobial production systems"

### Supplementary tables

**Table S1. Base case model parameters**

| Parameter | Value | Units | References |
|---|---|---|---|
| Operating conditions | | | |
| $P_{CO_2}$ | 0.4 | atm | -- |
| $D_{gas}$ | 100 | $hr^{-1}$ | -- |
| $T$ | 30 (*C. necator*) | °C | DSMZ |
| | 35 (*S. ovata*) | | DSMZ |
| | 37 (*E. coli*) | | DSMZ |
| Microbial growth | | | |
| *C. necator* (formatotrophy) | | | |
| $\mu_{max,opt}$ | 0.18 | $hr^{-1}$ | [1] |
| $Y'_{X/F,maz}$ | 0.169 | mol $mol^{-1}$ | [1] |
| $Y'_{L/F,maz}$ | 0.11 | mol $mol^{-1}$ | calculated |
| $\theta_F$ | 75.11 | mM | [1] |
| $K_{S,F}$ | 10 | μM | [2] |
| $K_{S,O_2}$ | 2.5 | μM | [3] |
| $pH_{opt}$ | 7 | -- | [1] |
| $pH_{min}$ | 4 | -- | [4] |
| $pH_{max}$ | 9 | -- | [4] |
| *C. necator* (hydrogenotrophy) | | | |
| $\mu_{max,opt}$ | 0.18 | $hr^{-1}$ | [5] |
| $Y'_{X/H_2}$ | 0.19 | mol $mol^{-1}$ | [5] |
| $Y'_{L/H_2}$ | 0.11 | mol $mol^{-1}$ | calculated |
| $K_{S,H_2}$ | 20.4 | μM | [6] |
| $K_{S,O_2}$ | 2.5 | μM | [3] |
| $K_{S,CO_2}$ | 9.38 | μM | [6] |
| $pH_{opt}$ | 7 | -- | [1] |
| $pH_{min}$ | 4 | -- | [4] |
| $pH_{max}$ | 9 | -- | [4] |

| | | | |
|---|---|---|---|
| *S. ovata* (acetogenesis) | | | |
| $\mu_{\mathrm{max,opt}}$ | 0.044 | $hr^{-1}$ | [7] |
| $K_{\mathrm{S,H_2}}$ | 20 | μM | [8] |
| $K_{\mathrm{S,CO_2}}$ | 20 | μM | [8] |
| $\eta_{\mathrm{B}}$ | 0.6 | -- | [9] |
| $\mathrm{pH_{opt}}$ | 7 | -- | [7] |
| $\mathrm{pH_{min}}$ | 4 | -- | assumed |
| $\mathrm{pH_{max}}$ | 9 | -- | assumed |
| *E. coli* (acetotrophy) | | | |
| $\mu_{\mathrm{max,opt}}$ | 0.3 | $hr^{-1}$ | [10] |
| $Y'_{\mathrm{X/Ac}}$ | 0.936 | mol $mol^{-1}$ | [10] |
| $Y'_{\mathrm{L/Ac}}$ | 0.5 | mol $mol^{-1}$ | calculated |
| $K_{\mathrm{S,O_2}}$ | 2.5 | μM | [3] |
| $K_{\mathrm{S,Ac}}$ | 10 | μM | assumed |
| $K_{\mathrm{I,Ac}}$ | 0.83 | M | [11] |
| $\mathrm{pH_{opt}}$ | 7 | -- | [10] |
| $\mathrm{pH_{min}}$ | 4 | -- | [12–14] |
| $\mathrm{pH_{max}}$ | 9.5 | -- | [12–14] |
| Acid/base reactions | | | |
| $S_1$ | -96.31 | J $mol^{-1}$ $K^{-1}$ | [15] |
| $S_2$ | -148.1 | J $mol^{-1}$ $K^{-1}$ | [15] |
| $S_5$ | -71.0 | J $mol^{-1}$ $K^{-1}$ | [15] |
| $S_6$ | -92.4 | J $mol^{-1}$ $K^{-1}$ | [15] |
| $S_{\mathrm{w}}$ | -80.66 | J $mol^{-1}$ $K^{-1}$ | [15] |
| $H_1$ | 7.64 | kJ $mol^{-1}$ | [15] |
| $H_2$ | 14.85 | kJ $mol^{-1}$ | [15] |
| $H_5$ | -0.12 | kJ $mol^{-1}$ | [15] |
| $H_6$ | -0.4 | kJ $mol^{-1}$ | [15] |
| $H_{\mathrm{w}}$ | 55.84 | kJ $mol^{-1}$ | [15] |
| $K_7$ | $1.38 \times 10^{-4}$ | mol $L^{-1}$ | [15] |
| $k_{+1}$ | $\exp\left(1246.98 - \frac{6\times10^4}{T} - 183\ln(T)\right)$ | $s^{-1}$ | [16] |
| $k_{+2}$ | 59.44 | $s^{-1}$ | [16] |
| $k_{+3}$ | $2.23 \times 10^3$ | L $mol^{-1}$ $s^{-1}$ | [16] |
| $k_{+4}$ | $6.0 \times 10^9$ | L $mol^{-1}$ $s^{-1}$ | [16] |
| $k_{+5}$ | 10 | $s^{-1}$ | assumed |
| $k_{+6}$ | 10 | $s^{-1}$ | assumed |
| $k_{+7}$ | 10 | $s^{-1}$ | assumed |
| $k_{+\mathrm{w}}$ | $2.4 \times 10^{-5}$ | L $mol^{-1}$ $s^{-1}$ | [17] |
| Gas/liquid mass transfer | | | |
| $k_{\mathrm{L}}a_{\mathrm{O_2}}$ | 200 | $hr^{-1}$ | assumed |
| $A_{\mathrm{S}}$ | 0.56 | $m^{-1}$ | assumed |
| Diffusion coefficients | | | |
| $D_{\mathrm{CO_2}}$ | $14.68 \times 10^{-9}\left(\frac{T}{217.206} - 1\right)^{1.997}$ | $m^2$ $s^{-1}$ | [18] |
| $D_{\mathrm{H_2}}$ | $\frac{2.290\times10^{-11}}{\mu^{0.819}}T$ | $m^2$ $s^{-1}$ | [2] |
| $D_{\mathrm{O_2}}$ | $10\hat{}\left[-8.410 + \frac{773.8}{T} - \left(\frac{506.4}{T}\right)^2\right]$ | $m^2$ $s^{-1}$ | [19] |
| Bunsen coefficients | | | |

| | | | |
|---|---|---|---|
| $A_{1,CO_2}$ | -60.2409 | -- | [20] |
| $A_{2,CO_2}$ | 93.4517 | -- | [20] |
| $A_{3,CO_2}$ | 23.3585 | -- | [20] |
| $B_{1,CO_2}$ | $2.3517 \times 10^{-2}$ | -- | [20] |
| $B_{2,CO_2}$ | $-2.3656 \times 10^{-2}$ | -- | [20] |
| $B_{3,CO_2}$ | $4.7036 \times 10^{-3}$ | -- | [20] |
| $A_{1,O_2}$ | -58.3877 | -- | [21] |
| $A_{2,O_2}$ | 85.8079 | -- | [21] |
| $A_{3,O_2}$ | 23.8439 | -- | [21] |
| $B_{1,O_2}$ | $3.4892 \times 10^{-2}$ | -- | [21] |
| $B_{2,O_2}$ | $1.5568 \times 10^{-2}$ | -- | [21] |
| $B_{3,O_2}$ | $-1.9387 \times 10^{-3}$ | -- | [21] |
| $A_{1,H_2}$ | -39.9611 | -- | [22] |
| $A_{2,H_2}$ | 53.9381 | -- | [22] |
| $A_{3,H_2}$ | 16.3135 | -- | [22] |
| $B_{1,H_2}$ | $2.3517 \times 10^{-2}$ | -- | [22] |
| $B_{2,H_2}$ | $1.7566 \times 10^{-2}$ | -- | [22] |
| $B_{3,H_2}$ | $-2.3010 \times 10^{-3}$ | -- | [22] |
| pH controller | | | |
| $K_C$ | 0.1 | $hr^{-1}$ | -- |
| $\tau$ | 60 | s | -- |
| Combustion energy | | | |
| $\Delta_r G_X^0$ | -479 | $kJ\ mol^{-1}$ | Note 2 |
| $\Delta_r G_E^0$ | -479 | $kJ\ mol^{-1}$ | Note 2 |
| $\Delta_r G_{H_2}^0$ | -260 | $kJ\ mol^{-1}$ | Note 2 |
| $\Delta_r G_{FFA}^0$ | -240 | $kJ\ mol^{-1}$ | Note 2 |
| $\Delta_r G_{LLA}^0$ | -1370 | $kJ\ mol^{-1}$ | Note 2 |
| $\Delta_r G_{AAA}^0$ | -870 | $kJ\ mol^{-1}$ | Note 2 |
| $CO_2$ electrolyzer | | | |
| $j$ | 140 | $mA\ cm^{-2}$ | [23] |
| $\eta_F$ | 94 | % | [23] |
| $V_e$ | 3.5 | V | [23] |
| $c_{FFA,eff}$ | 2.08 | M | [23] |
| $H_2$ electrolyzer | | | |
| $j$ | 1000 | $mA\ cm^{-2}$ | [24] |
| $\eta_F$ | 99 | % | [24] |
| $V_e$ | 2.0 | V | [24] |

**Table S2. Parameter sensitivity**

| Parameter | Baseline Value | Units | Parameter -30% | Parameter +30% | Formate GWP -30% | H2 GWP -30% | Acetate GWP -30% | Hetero GWP -30% | Formate GWP +30% | H2 GWP +30% | Acetate GWP +30% | Hetero GWP +30% |
|---|---|---|---|---|---|---|---|---|---|---|---|---|
| **Impact Model Parameters** | | | | | | | | | | | | |
| BASELINE | n/a | | n/a | n/a | **1.1684** | **0.6884** | **1.2488** | **1.9304** | **1.1684** | **0.6884** | **1.2488** | **1.9304** |
| Glucose GWP | 0.95 | kg co2/kg | 0.665 | 1.235 | 1.1684 | 0.6884 | 1.2488 | 1.3604 | 1.1684 | 0.6884 | 1.2488 | 2.5004 |
| % GWP Corn Fertilizer | 0.25 | | 0.175 | 0.325 | 1.1684 | 0.6884 | 1.2488 | 2.0229 | 1.1684 | 0.6884 | 1.2488 | 1.8379 |
| Heterotroph Productivity | 35.2 | kg/L/yr | 24.64 | 45.76 | 1.1684 | 0.6884 | 1.2488 | 1.9367 | 1.1684 | 0.6884 | 1.2488 | 1.927 |
| Glucose Yield | 0.5 | kg biomass/kg glucose | 0.35 | 0.65 | 1.1684 | 0.6884 | 1.2488 | 2.6368 | 1.1684 | 0.6884 | 1.2488 | 1.5501 |
| Distance NH3 Shipped [km] | 500 | km | 350 | 650 | 1.1669 | 0.6869 | 1.2469 | 1.9289 | 1.1699 | 0.69 | 1.2506 | 1.9319 |
| Distance Nutrients Shipped [km] | 500 | km | 350 | 650 | 1.1657 | 0.6845 | 1.2232 | 1.9265 | 1.1712 | 0.6923 | 1.2743 | 1.9343 |
| Distance electrolyzer materials Shipped [km] | 500 | km | 350 | 650 | 1.1684 | 0.6884 | 1.2488 | 1.9304 | 1.1684 | 0.6884 | 1.2488 | 1.9304 |
| Distance Bioreactors shipped [km] | 500 | km | 350 | 650 | 1.1683 | 0.6884 | 1.2486 | 1.9304 | 1.1685 | 0.6885 | 1.2489 | 1.9304 |
| Distance Glucose shipped [km] | 500 | km | 350 | 650 | 1.1684 | 0.6884 | 1.2488 | 1.9134 | 1.1684 | 0.6884 | 1.2488 | 1.9474 |
| Wind GWP | 9.102 | kg co2/MWh | 6.3714 | 11.8326 | 1.0216 | 0.6099 | 1.1646 | 1.9217 | 1.3152 | 0.767 | 1.3329 | 1.9392 |
| DAC Electricity Demand | 2.02 | kWh/kg CO2 | 1.414 | 2.626 | 1.1578 | 0.6788 | 1.2379 | 1.9304 | 1.179 | 0.6981 | 1.2597 | 1.9304 |
| Adsorbent GWP | 23.7 | g CO2-e/kg CO2 capture | 16.59 | 30.81 | 1.1547 | 0.6759 | 1.2347 | 1.9304 | 1.1821 | 0.7009 | 1.2628 | 1.9304 |
| DAC Plant GWP | 15 | g CO2-e/kg CO2 capture | 10.5 | 19.5 | 1.1597 | 0.6805 | 1.2399 | 1.9304 | 1.1771 | 0.6963 | 1.2577 | 1.9304 |
| GWP Ammonia (besides H2 electrolysis) | 0.4105 | kg co2/ kg nh3 | 0.28735 | 0.53365 | 1.1464 | 0.6664 | 1.2224 | 1.8934 | 1.1904 | 0.7105 | 1.2751 | 1.9674 |
| Green Ammonia Electricity Demand | 9.59 | kWh/kg NH3 | 6.713 | 12.467 | 1.1638 | 0.6838 | 1.2433 | 1.9227 | 1.173 | 0.693 | 1.2543 | 1.9381 |
| Phosphoric Acid GWP | 1.012 | kg CO2-e/kg | 0.7084 | 1.3156 | 1.137 | 0.6571 | 1.2112 | 1.899 | 1.1998 | 0.7198 | 1.2863 | 1.9618 |
| NaCl GWP | 0.178 | kg CO2-e/kg | 0.1246 | 0.2314 | 1.1684 | 0.6884 | 1.2488 | 1.9304 | 1.1684 | 0.6884 | 1.2488 | 1.9304 |
| MgSO4 GWP | 0.239 | kg CO2-e/kg | 0.1673 | 0.3107 | 1.1654 | 0.6855 | 1.2452 | 1.9274 | 1.1714 | 0.6914 | 1.2523 | 1.9334 |
| CaCl2 GWP | 0.611 | kg CO2-e/kg | 0.4277 | 0.7943 | 1.1673 | 0.6873 | 1.2474 | 1.9293 | 1.1695 | 0.6896 | 1.2501 | 1.9315 |
| FeCl3 GWP | 0.511 | kg CO2-e/kg | 0.3577 | 0.6643 | 1.1684 | 0.6884 | 1.2487 | 1.9304 | 1.1685 | 0.6885 | 1.2488 | 1.9305 |
| NaOH GWP minus electrolysis | 0.116 | kg CO2-e/kg | 0.0812 | 0.1508 | 1.1684 | 0.6884 | 1.204 | 1.9304 | 1.1684 | 0.6884 | 1.2935 | 1.9304 |
| HCl GWP minus electrolysis | 0.127 | kg CO2-e/kg | 0.0889 | 0.1651 | 1.1618 | 0.6767 | 1.1901 | 1.9187 | 1.175 | 0.7001 | 1.3074 | 1.9421 |
| NaOH chlor-alkili electricity demand | 1.1 | kWh/kg | 0.77 | 1.43 | 1.1684 | 0.6884 | 1.2449 | 1.9304 | 1.1684 | 0.6884 | 1.2526 | 1.9304 |

| | | | | | | | | | | | | |
|---|---|---|---|---|---|---|---|---|---|---|---|---|
| HCl Chlor-alkili electricity demand | 1.21 | kWh/kg | 0.847 | 1.573 | 1.1678 | 0.6874 | 1.2437 | 1.9294 | 1.169 | 0.6894 | 1.2538 | 1.9314 |
| Iridium GWP | 8860 | kg Co2-e/kg | 6202 | 11518 | 1.1468 | 0.6869 | 1.2474 | 1.9304 | 1.19 | 0.69 | 1.2501 | 1.9304 |
| Carbon Fibre GWP | 30.98 | kg Co2-e/kg | 21.686 | 40.274 | 1.1675 | 0.6883 | 1.2487 | 1.9304 | 1.1694 | 0.6885 | 1.2489 | 1.9304 |
| Tin GWP | 5.77 | kg Co2-e/kg | 4.039 | 7.501 | 1.1684 | 0.6884 | 1.2488 | 1.9304 | 1.1684 | 0.6884 | 1.2488 | 1.9304 |
| Nafion GWP | 831 | kg Co2-e/kg | 581.7 | 1080.3 | 1.1118 | 0.6872 | 1.2477 | 1.9304 | 1.225 | 0.6896 | 1.2498 | 1.9304 |
| Polystyrene Sulfonate Cationic Resin GWP | 1.76 | kg Co2-e/kg | 1.232 | 2.288 | 1.168 | 0.6884 | 1.2488 | 1.9304 | 1.1688 | 0.6884 | 1.2488 | 1.9304 |
| Reactor Volume | 30000 | L | 21000 | 39000 | 1.1744 | 0.6938 | 1.2604 | 1.9312 | 1.1645 | 0.6849 | 1.241 | 1.9299 |
| Reactor Weight:Vol | 0.6235 | kg/L | 0.43645 | 0.81055 | 1.1543 | 0.6757 | 1.2211 | 1.9285 | 1.1826 | 0.7012 | 1.2764 | 1.9323 |
| Reactor Lifetime | 8 | yr | 5.6 | 10.4 | 1.1886 | 0.7066 | 1.2883 | 1.9331 | 1.1575 | 0.6786 | 1.2275 | 1.929 |
| Steel GWP | 6.89 | kg CO2-e/kg | 4.823 | 8.957 | 1.1543 | 0.6758 | 1.2212 | 1.9286 | 1.1825 | 0.7011 | 1.2763 | 1.9323 |
| Reactor Area Density | 50 | L/m^2 | 35 | 65 | 1.196 | 0.7132 | 1.3026 | 1.934 | 1.1536 | 0.6751 | 1.2197 | 1.9285 |
| Plant Lifetime | 40 | year | 28 | 52 | 1.196 | 0.7132 | 1.3026 | 1.934 | 1.1536 | 0.6751 | 1.2197 | 1.9285 |
| Bulding GWP/m^2 | 595.9 | kg CO2-e/kg | 417.13 | 774.67 | 1.1491 | 0.6711 | 1.211 | 1.9279 | 1.1877 | 0.7058 | 1.2865 | 1.933 |
| Nitrogen Ratio | 0.14 | g N/ g biomass | 0.098 | 0.182 | 1.1211 | 0.6411 | 1.2384 | 1.8831 | 1.2157 | 0.7358 | 1.2591 | 1.9777 |
| Phosphorous Ratio | 0.031 | g P/g biomass | 0.0217 | 0.0403 | 1.14 | 0.66 | 1.2055 | 1.902 | 1.1968 | 0.7168 | 1.292 | 1.9588 |
| Sulfur Ratio | 0.0105 | g S/g biomass | 0.00735 | 0.01365 | 1.1651 | 0.6851 | 1.2448 | 1.9271 | 1.1717 | 0.6917 | 1.2527 | 1.9337 |
| Calcium Ratio | 0.0021 | g Ca/g biomass | 0.00147 | 0.00273 | 1.1672 | 0.6873 | 1.2474 | 1.9292 | 1.1696 | 0.6896 | 1.2502 | 1.9316 |
| Iron Ratio | 0.0001 | g Fe/g biomass | 0.00007 | 0.00013 | 1.1684 | 0.6884 | 1.2487 | 1.9304 | 1.1685 | 0.6885 | 1.2488 | 1.9305 |
| Nitrient Utilization Ratio | 0.95 | | 0.665 | 1 ** | 1.2831 | 0.8031 | 1.3332 | 2.0451 | 1.155 | 0.6751 | 1.2389 | 1.917 |
| **Bioreactor Parameters** | | | | | | | | | | | | |
| BASELINE | n/a | | n/a | n/a | **1.1685** | **0.6884** | **1.2366** | **1.9304** | **1.1685** | **0.6884** | **1.2366** | **1.9304** |
| mu_max,opt | 0.18 | hr^-1 | 0.126 | 0.234 | 1.1685 | 0.6884 | 1.2366 | 1.9304 | 1.1685 | 0.6884 | 1.2366 | 1.9304 |
| Y' X,F max | 0.169 | mol/mol | 0.1183 | 0.2197 | 1.5374 | 0.6884 | 1.2366 | 1.9304 | 0.9713 | 0.6884 | 1.2366 | 1.9304 |
| theta_F | 75.11 | mM | 52.577 | 97.643 | 1.1685 | 0.6884 | 1.2366 | 1.9304 | 1.1685 | 0.6884 | 1.2366 | 1.9304 |
| Ks,F | 10 | uM | 7 | 13 | 1.1685 | 0.6884 | 1.2366 | 1.9304 | 1.1686 | 0.6884 | 1.2366 | 1.9304 |
| KS,O2 | 2.5 | uM | 1.75 | 3.25 | 1.1685 | 0.6884 | 1.2366 | 1.9304 | 1.1686 | 0.6884 | 1.2366 | 1.9304 |
| pHmin | 4 | | 2.8 | 5.2 | 1.1685 | 0.6884 | 1.2366 | 1.9304 | 1.1685 | 0.6884 | 1.2366 | 1.9304 |
| pHmax** | 9 | | 8 | 11.7 | 1.1685 | 0.6884 | 1.2366 | 1.9304 | 1.1685 | 0.6884 | 1.2366 | 1.9304 |
| mu_max,opt | 0.18 | hr^-1 | 0.126 | 0.234 | 1.1685 | 0.6881 | 1.2366 | 1.9304 | 1.1685 | 0.6887 | 1.2366 | 1.9304 |
| Y' X,H max | 0.19 | mol/mol | 0.133 | 0.247 | 1.1685 | 0.848 | 1.2366 | 1.9304 | 1.1685 | 0.6144 | 1.2366 | 1.9304 |

| | | | | | | | | | | | | |
|---|---|---|---|---|---|---|---|---|---|---|---|---|
| Ks,H2 | 20.4 | uM | 14.28 | 26.52 | 1.1685 | 0.6883 | 1.2366 | 1.9304 | 1.1685 | 0.6886 | 1.2366 | 1.9304 |
| Ks,O2 | 2.5 | uM | 1.75 | 3.25 | 1.1685 | 0.6884 | 1.2366 | 1.9304 | 1.1685 | 0.6885 | 1.2366 | 1.9304 |
| Ks,CO2 | 9.38 | uM | 6.566 | 12.194 | 1.1685 | 0.6884 | 1.2366 | 1.9304 | 1.1685 | 0.6884 | 1.2366 | 1.9304 |
| pHmin | 4 | | 2.8 | 5.2 | 1.1685 | 0.6884 | 1.2366 | 1.9304 | 1.1685 | 0.6884 | 1.2366 | 1.9304 |
| pHmax** | 9 | | 8 | 11.7 | 1.1685 | 0.6884 | 1.2366 | 1.9304 | 1.1685 | 0.6884 | 1.2366 | 1.9304 |
| mu_max_opt | 0.044 | hr^-1 | 0.0308 | 0.0572 | 1.1685 | 0.6884 | 1.2998 | 1.9304 | 1.1685 | 0.6884 | 1.202 | 1.9304 |
| Ks,H2 | 20 | uM | 14 | 26 | 1.1685 | 0.6884 | 1.2331 | 1.9304 | 1.1685 | 0.6884 | 1.2387 | 1.9304 |
| Ks,CO2 | 20 | uM | 14 | 26 | 1.1685 | 0.6884 | 1.2366 | 1.9304 | 1.1685 | 0.6884 | 1.2366 | 1.9304 |
| eta_B | 0.6 | | 0.42 | 0.78 | 1.1685 | 0.6884 | 1.2229 | 1.9304 | 1.1685 | 0.6884 | 1.2489 | 1.9304 |
| pHmin | 4 | | 2.8 | 5.2 | 1.1685 | 0.6884 | 1.2366 | 1.9304 | 1.1685 | 0.6884 | 1.2366 | 1.9304 |
| pHmax** | 9 | | 8 | 11.7 | 1.1685 | 0.6884 | 1.2366 | 1.9304 | 1.1685 | 0.6884 | 1.2366 | 1.9304 |
| mu_max_opt | 0.3 | hr^-1 | 0.21 | 0.39 | 1.1685 | 0.6884 | 1.2366 | 1.9304 | 1.1685 | 0.6884 | 1.2365 | 1.9304 |
| Y' X,Ac | 0.936 | mol/mol | 0.6552 | 1.2168 | 1.1685 | 0.6884 | 1.6882 | 1.9304 | 1.1685 | 0.6884 | 0.9996 | 1.9304 |
| Ks,O2 | 2.5 | uM | 1.75 | 3.25 | 1.1685 | 0.6884 | 1.2366 | 1.9304 | 1.1685 | 0.6884 | 1.2366 | 1.9304 |
| Ks,Ac | 10 | uM | 7 | 13 | 1.1685 | 0.6884 | 1.2366 | 1.9304 | 1.1685 | 0.6884 | 1.2366 | 1.9304 |
| Ki,Ac | 0.83 | uM | 0.581 | 1.079 | 1.1685 | 0.6884 | 1.2366 | 1.9304 | 1.1685 | 0.6884 | 1.2366 | 1.9304 |
| pHmin | 4 | | 2.8 | 5.2 | 1.1685 | 0.6884 | 1.2366 | 1.9304 | 1.1685 | 0.6884 | 1.2366 | 1.9304 |
| pHmax** | 9.5 | | 8 | 12.35 | 1.1685 | 0.6884 | 1.2366 | 1.9304 | 1.1685 | 0.6884 | 1.2366 | 1.9304 |

**Supplementary notes**

*Note 1: Polylactic Acid Production Life Cycle Impacts*

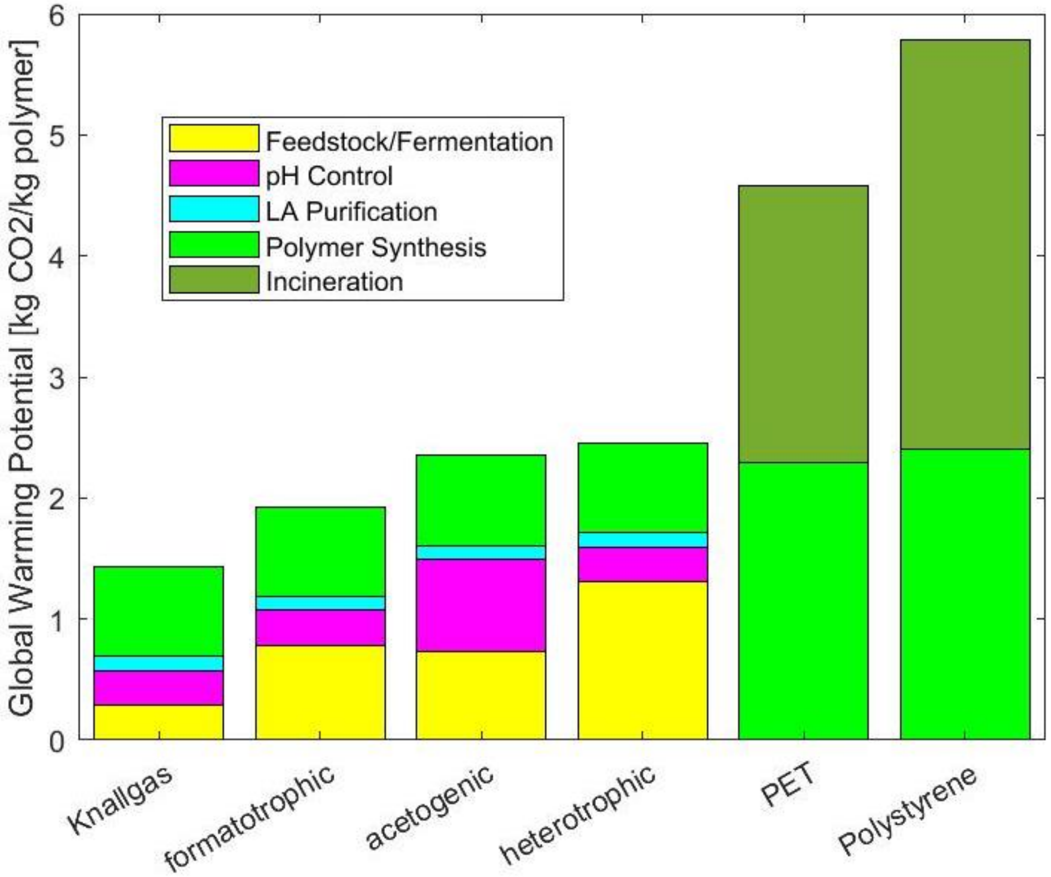

**Figure S1: Life Cycle Global Warming Potential of EMP-based and traditional plastics.** Life cycle (cradle-to-grave) global warming potential of polymer production of polylactic acid (PLA) in the three EMP systems and traditional bioprocesses compared to those of fossil-fuel based plastics polyethylene terephthalate (PET) and polystyrene (PS). EMP production assumes a 90% carbon efficiency (as defined in main text) and grid composed solely of wind power.

We analyzed the cradle-to-grave life cycle impact of polylactic acid (PLA) production. We split the PLA production process into five categories: feedstock production and fermentation, pH control, lactic acid purification, polymer synthesis, and end of life. The global warming potentials for the lactic acid production and pH control are identical to the results in Fig. 4c in the main text. Purification of lactic acid relies on acidifying the lactate anion produced in the bioreactor and therefore requires a stoichiometric proportion of sulfuric acid. We obtained data for the carbon footprint of polylactic acid polymer production from Morão and de Bie.[25] We further compared the global warming footprints to those of two major fossil-fuel based polymers: polyethylene terephthalate (PET) and polystyrene (PS). Carbon footprints for the production of these two polymers was obtained from the PEF dataset. Two end-of-life scenarios were considered: burial and incineration. In the case of burial, no carbon dioxide is released. In the case of incineration, the carbon footprint is equal to the stoichiometric amount of carbon dioxide that would be produced by complete combustion of the polymer.

The global warming potentials of the cradle-to-grave PLA production of the three EMP systems and the traditional bioprocess reflect the same trends as for lactic acid production because the lactic acid purification and polymer synthesis steps are identical for each system. Therefore, the Knallgas bacteria-based system outcompetes the other bioprocesses in PLA production as it does in lactic acid production. The $H_2$-mediated and formate-mediated systems also clearly outperforms the two fossil-based polymers, PET and PS, in terms of life cycle global warming potential, in the scenario shown. The acetogenic system and heterotrophic system both have comparable carbon footprints to the fossil-based plastics if the plastics are not incinerated. However, a true like-to-like comparison will involve incinerating these plastics to leave no waste, as PLA will ultimately biodegrade in the environment. In the scenario, all methods of PLA production have smaller carbon footprints when compared to PET and PS.

These results are for a carbon conversion efficiency of 90%, similar to current glucose-based PLA fermentation. As noted in the main text, an efficiency this high will be difficult in EMP processes. Lower carbon efficiencies in the lactic acid production step will therefore increase the total carbon footprint of PLA production. As shown in Fig. 5b in the main text, this change in GWP is also dependent on the electricity source/grid composition. As with lactic acid production, for $H_2$-mediated electromicrobial production of PLA to outcompete heterotroph-based processes, the process must rely on renewable electricity and must achieve a carbon efficiency of around 50%.

*Note 2: calculating combustion energies*

To calculate combustion energies, we adopted the strategy of Claassens *et al.*[26] Briefly, we used eQuilibrator[27] to calculate the $\Delta_{\mathrm{r}}G'^{0}$ of the combustion reaction at a pH of 7.0 and ionic strength of 0.1 M. The biomass combustion energy was adopted from previous calculations.[28,29]

*Note 3: reactor model parameter sensitivity calculations*

When some parameters (*e.g.* the maximum specific growth rate, $\mu_{\mathrm{max,opt}}$) are adjusted, operating conditions such as the liquid dilution rate ($D_{\mathrm{liq}}$) have different optima. In these cases, we identified the new optimal operating conditions and report system outputs based on these.
